# Retrospective harmonization of multi-site diffusion MRI data acquired with different acquisition parameters

**DOI:** 10.1101/314179

**Authors:** Suheyla Cetin Karayumak, Sylvain Bouix, Lipeng Ning, Martha Shenton, Marek Kubicki, Yogesh Rathi

**Author notes:** Corresponding author. Email address (Suheyla Cetin Karayumak).

## Abstract

A joint and integrated analysis of multi-site diffusion MRI (dMRI) datasets can dramatically increase the statistical power of neuroimaging studies and enable comparative studies pertaining to several brain disorders. However, dMRI data sets acquired on multiple scanners cannot be naively pooled for joint analysis due to scanner specific nonlinear effects as well as differences in acquisition parameters. Consequently, for joint analysis, the dMRI data has to be harmonized, which involves removing scanner-specific differences from the raw dMRI signal. In this work, we present a dMRI harmonization method that, when applied to multi-site data, is capable of removing scanner-specific effects, while accounting for minor differences in acquisition parameters such as b-value, spatial resolution and number of gradient directions in the dMRI data (typical for multi-site clinical research scans). We validate our algorithm on dMRI data acquired from two sites: Philadelphia Neurodevelopmental Cohort (PNC) with 800 healthy adolescents (ages 8 to 22 years) and Brigham and Women’s Hospital (BWH) with 70 healthy subjects (ages 14 to 54 years). In particular, we show that gender differences and maturation in different age groups are preserved after harmonization, as measured using effect sizes (small, medium and large), irrespective of the test sample size. Further, because we use matched control subjects from different scanners to estimate scanner-specific effects, we tested how many subjects are needed from each site to achieve best harmonization results. Our results indicate that at-least 16 to 18 well-matched healthy controls from each site are needed to reliably capture scanner related differences. The proposed method can thus be used for retrospective harmonization of raw dMRI data across sites despite differences in acquisition parameters, while preserving inter-subject anatomical variability.

## 1. Introduction

Diffusion MRI is sensitive to molecular water motion, which can be recorded non-invasively by an MRI scanner. However, these measurements are affected by different hardware specifications (magnetic field strength, number of the receiver coils etc.), and different acquisition parameters (echo time, diffusion time, gradient strength, voxel size, number of gradient directions etc.) (Helmer et al., 2016). Therefore, the data acquired by each scanner is substantially different even for the same subject. In fact, even if the same subject is scanned with the same hardware from the same manufacturer, diffusion signal can still be different (Vollmar et al., 2010). This is due to differences in magnetic field inhomogeneities, sensitivity of receiver coils, the number of receiver coils used, vendor-specific MRI reconstruction algorithms and differences in acquisition parameters. Consequently, dMRI data must be harmonized prior to joint analysis.

Several methods have characterized both intra-scanner and inter-scanner variability in structural and dMRI data (Landman et al., 2011, 2007). Based on their study in Walker et al. (2013), the authors recommend the use of physical phantoms to monitor and quickly detect any scanner-related changes in ongoing neuroimaging studies. While the use of physical phantoms is necessary, they are inadequate in capturing the regional and tissue specific scanner differences. Further, it is non-trivial to use the scanner differences observed in physical phantoms to correct human in-vivo data, due to the complexities of biological tissue.

Existing techniques on data pooling are based on using diffusion tensor imaging (DTI) derived metrics (Salimi-Khorshidi et al., 2009; Jahanshad et al., 2013; Kochunov et al., 2014; Forsyth et al., 2014; Venkatraman et al., 2015; Jenkins et al., 2016; Pohl et al., 2016; Fortin et al., 2017). For instance, Salimi-Khorshidi et al. (2009); Jahanshad et al. (2013); Kochunov et al. (2014); Palacios et al. (2016); Kelly et al. (2017)use meta-analysis approach which involves combining z-scores of a given diffusion measure (e.g. fractional anisotropy (FA)) from all sites to determine group differences. However, the subject population at each site may not be sufficient to capture the variance of the entire population, a critical requirement to ensure proper pooling and analysis of the z-scores (which depends on the variance and not just the population mean). Further, z-scores may not be the best statistic to use if the distribution of the diffusion measure in the population is not Gaussian (normal). On the other hand, Forsyth et al. (2014); Venkatraman et al. (2015); Fortin et al. (2017) use statistical covariates to regress out the differences between sites in DTI measures such as FA, mean diffusivity (MD) or cortical thickness. Of particular note is the work of Pohl et al. (2016), where the authors use information from 3 traveling subjects to obtain a linear correction factor for scanner related effects in FA (a different correction factor for each ROI analyzed). This method however has limitations when using large ROIs (such as the corticospinal tract), as the scanner-related effects are not only non-linear but also regionally varying (see (Mirzaalian et al., 2016) and Figure 3). Thus, due to the regional variability of the diffusion signal, using a single regressor for large ROIs can lead to erroneous results in the aggregated data (Mirzaalian et al., 2016; Fortin et al., 2017).

**Figure 1:**
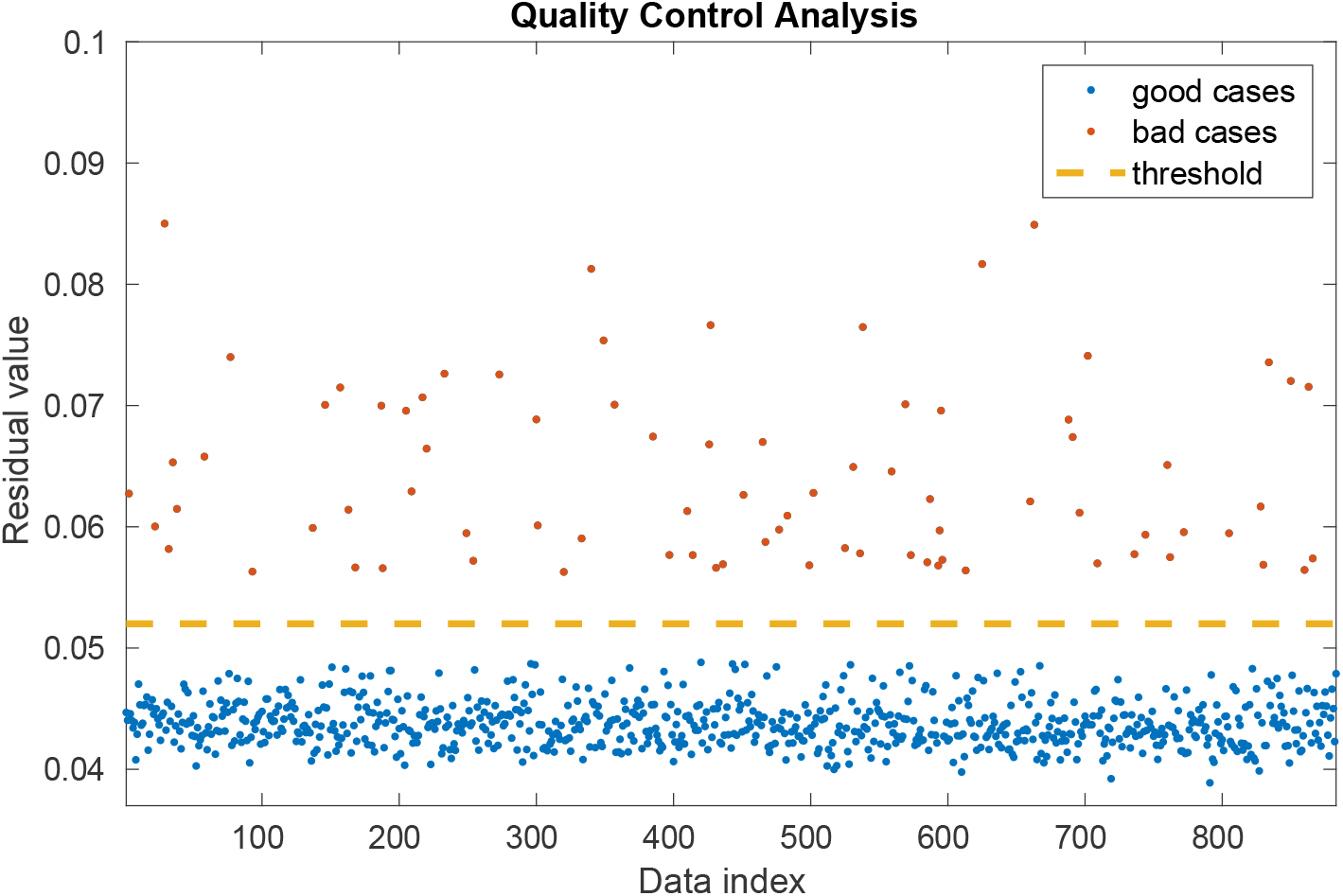
PNC data automated quality control analysis results: The average signal residual for each subject (over the entire brain) was calculated and this generated two clusters: for bad quality (orange) and for good quality (blue) cases. The threshold (yellow) to separate good and bad clusters was chosen in a heuristic manner.

**Figure 2:**
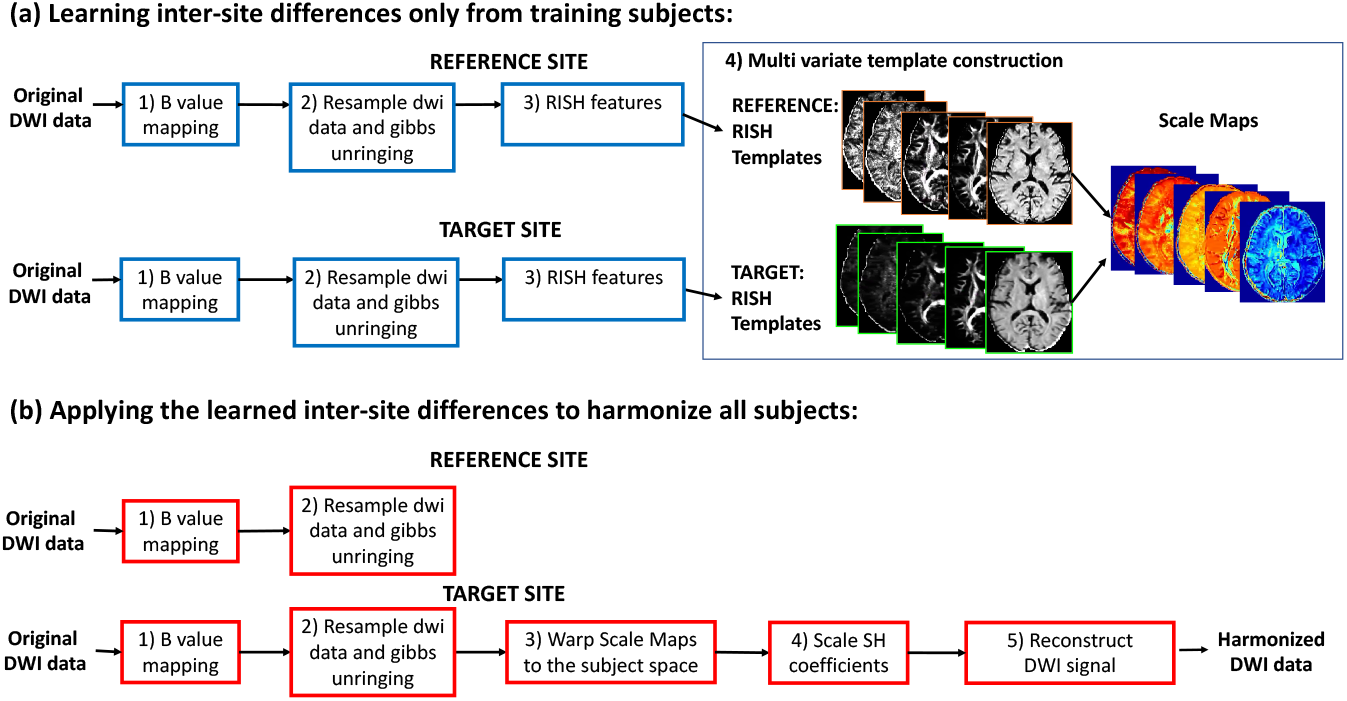
Steps of multi-variate RISH feature template construction and dMRI data harmonization.

**Figure 3:**
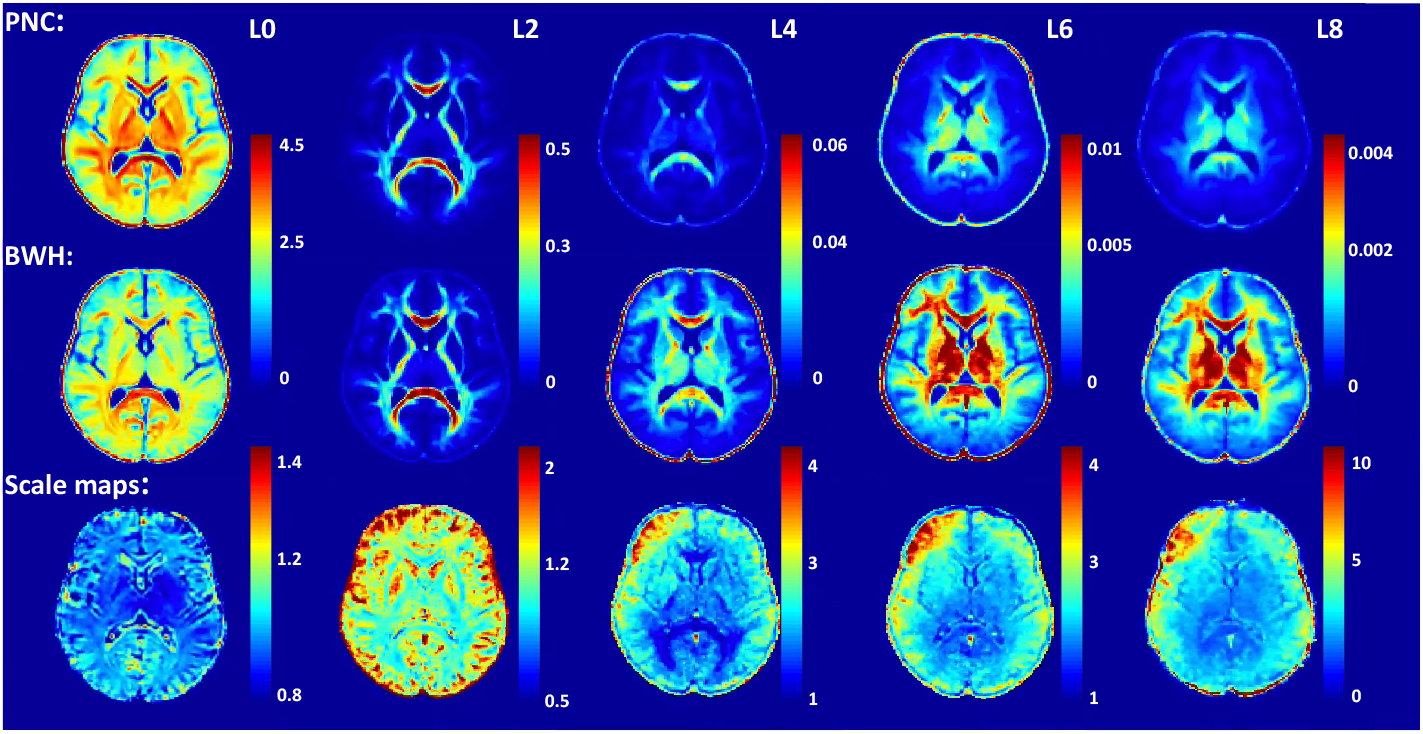
RISH Features for SH orders of *l* = {0, 2, 4, 6, 8} are depicted in each sub-figure from left to right for PNC site (top row) and BWH site (middle row). Scale maps for each RISH feature show the between-site mapping obtained between controls from two sites (bottom row).

All of the methods mentioned above have to correct for scanner-specific effects in each diffusion measure of interest separately, i.e., a linear correction factor for each diffusion measure, thus making the harmonization procedure entirely model-specific (e.g. single tensor). Recently, Fortin et al. (2017) have proposed a retrospective multi-site harmonization method that uses Combat (a batch-effect correction tool used in genomics) to remove the site effects from FA and MD. This method estimates an additive and a multiplicative site-effect coefficient at each voxel, thus accounting for regional scanner differences. Despite this, their optimization procedure assumes that the site-effect parameters follow a particular parametric prior distribution (Gaussian and Inverse-gamma), which might not generalize to all scenarios or measures derived from other models (e.g., multi-compartment models).

### Contributions of this work

In our earlier works (Mirzaalian et al., 2016, 2017), we had proposed a model-free dMRI harmonization method which can be used to harmonize the “raw dMRI signal” (and not just a particular dMRI measure of interest) across sites. However, that work exclusively focused on harmonizing dMRI data across sites but with similar acquisition parameters. Thus, the method worked only when the spatial resolution and b-values were the same across sites. Additionally, the earlier method did not have an extensive validation on a large dataset.

In this work, we further build on our existing framework and propose a model free harmonization method that learns an efficient mapping across scanners despite differences in scanner parameters. We extensively validate our algorithm on dMRI data acquired from two different sites with different acquisition parameters. We use two independent data sets of different sizes (BWH: 70 subjects and PNC: 800 subjects) to demonstrate that our harmonization method is not affected by the sample size as opposed to existing approaches that require an accurate estimate of the variance of the underlying population in their model (e.g. meta-analysis methods). To this end, we compute effect sizes between groups separated by age and sex. Specifically, we show that the effect sizes, whether small, medium or large, are preserved by our harmonization procedure in both small (e.g. BWH) and large (e.g. PNC) data sets. Such validation experiments are necessary to robustly demonstrate the generalizability of any harmonization procedure for use with clinical research dMRI studies. Most importantly, using bootstrap experiments, we find that at-least 16 to 18 well-matched subjects per site are needed to robustly learn the mapping between sites.

## 2. Methods

### 2.1. Data Collection and Preprocessing

#### Neurodevelopmental Cohort (PNC)

We used dMRI data from 884 healthy participants from the publically available NIH repository: Philadelphia Neurodevelopmental Cohort (PNC) study (Satterthwaite et al., 2014, 2016). The dMRI data was acquired on a Siemens TIM Trio whole-body scanner, using a 32 channel head coil and a twice-refocused spin-echo (TRSE) single-shot EPI sequence with the following parameters: *TR* = 8100*ms* and *TE* = 82*ms*, b-value of 1000*s/mm*^2^, 7 *b* = 0 images. DMRI data was acquired with 64 diffusion-weighted directions divided into two independent sets, each with 32 diffusion-weighted directions. The images were acquired at 1.875 × 1.875 × 2 *mm*^3^ spatial resolution.

#### Brigham and Women’s Hospital (BWH)

DMRI images from healthy volunteers were acquired on a whole body General Electric MRI scanner (GE Medical Systems, Milwaukee) at Brigham and Women’s Hospital as part of a larger NIH funded study. A high resolution diffusion acquisition with the following parameters was used: twice refocused, *TR* = 17*s*, *TE* = 80*ms*, 1.67 × 1.67 × 1.7*mm*^3^ spatial resolution, 51 gradient directions with *b* = 900*s/mm*^2^ and eight additional *b* = 0 images.

Table 1 summarizes the demographic information for each of these sites. We applied axis alignment, centering and eddy current correction to each acquisition separately using the Psychiatry Neuroimaging Laboratory (PNL) pipeline: https://github.com/pnlbwh/pnlutil. We used the brain extraction tool (BET) to generate the brain masks (Smith, 2002; Jenkinson et al., 2005). The two dMRI acquisitions in the PNC data were combined by registering their respective baselines using affine transformation (Advanced Normalization Tools (ANTs) (Avants et al., 2011)). Then, the transformation was applied to each diffusion weighted volume and the gradient vectors were rotated using the rotation matrix estimated from the affine transformation. After merging the acquisitions, we performed an automated quality check of all 884 PNC data sets as follows: We fit the dMRI signal at each voxel using spherical harmonic basis functions (up to 8*^th^* order) (Descoteaux et al., 2007). Next, the average signal residual for each subject (over the entire brain) was calculated. This produced two clusters, one affected by motion and signal drops (bad cases) and another for good quality cases. We removed the cases with highest average residual, categorized as bad quality cases (84 participants in total). The threshold to determine the bad cases was manually chosen to maximize the separation between the clusters (see Figure 1).

**Table 1:** Demographics and dMRI acquisition information of the studies and harmonized results.

| Dataset | # Sub | Age | Gender | IQ | Handedness | dMRI data |
| --- | --- | --- | --- | --- | --- | --- |
| PNC | 800 | 8 to 22 years<br>(15.12±3.26) | 420 F<br>380 M | 101.89±16.33 | 12.5% L<br>87.5% R | b=1000 s/mm<sup>2</sup> |
|  |  |  |  |  |  | 64 directions |
|  |  |  |  |  |  | TE/TR=82/8100 ms<br>resolution=1.875 × 1.875 × 2mm<sup>3</sup> |
| BWH | 70 | 14 to 54 years<br>(30.34±12.35) | 22 F<br>48 M | 114.71±14.68 | 8% L<br>92% R | b=900 s/mm<sup>2</sup> |
|  |  |  |  |  |  | 51 directions |
|  |  |  |  |  |  | TE/TR=80/17000 ms<br>resolution=1.67 × 1.67 × 1.7mm<sup>3</sup> |
| Harmonized | 870 | 8 to 54 years<br>(16.43±6.40) | 412 F<br>458 M | 101.96±16.36 | 12.5% L<br>87.5% R | b=1000 s/mm<sup>2</sup> |
|  |  |  |  |  |  | resolution=1.5 × 1.5 × 1.5mm<sup>3</sup> |

*Abbreviations:* Dataset: PNC - Philadelphia Neurodevelopmental Cohort; BWH - Brigham and Women’s Hospital; F: females; M: males; R: right-handed, L: left-handed.

BWH data was also processed using the same PNL pipeline. Since the sample size is smaller, it was manually inspected for any signal dropouts or artifacts (as part of a separate study) and all subjects who did not pass our quality control procedure were not included in this study. A total of 70 subjects were included in this study after quality control analysis. See Table 1 for demographics of both the PNC and BWH data.

### 2.2. Group matching of training subjects across sites

Initially, we selected 20 right-handed (10F+10M) subjects from each site (detailed analysis related to the training subjects size is going to be explained in Section 2.4). The subjects were matched across sites for age and IQ to the best possible extent using unpaired t-test to minimize the statistical biological differences across sites. See Table 2 for demographics of training data. These training subjects were then used to learn the scanner-specific differences between sites. Details about the harmonization procedure is explained in the subsequent sections.

**Table 2:** Demographics of training subjects at PNC and BWH sites.

| Dataset | # Sub | Age | Gender | IQ | Handedness |
| --- | --- | --- | --- | --- | --- |
| PNC | 20 | 15 to 23 years | 10 F | 110.89±6.33 | 20 R |
|  |  | (19.37±2.19) | 10 M |  |  |
| BWH | 20 | 14 to 23 years | 10 F | 110.23±6.27 | 20 R |
|  |  | (19.57±2.26) | 10 M |  |  |

### 2.3. Steps for voxel-wise harmonization

The overall outline of the proposed method is depicted in a flowchart in Figure 2. Briefly, we first project the signal from both sites to a common canonical space of b-values and spatial resolution. Next, a set of matched controls are used to learn a non-linear mapping (in the dMRI signal domain) that captures scanner-specific differences between the sites (see Figure 2a). This mapping is then used to update the dMRI signal for each subject at the target site (see Figure 2b), i.e., we harmonize the remaining set of subjects from the target site. Each step in this process is explained in detail in the following subsections.

#### 2.3.1. B-value mapping and resampling

Due to differences in the b-values between sites, we first match the b-values for both the sites. Using evidence from existing works (Jensen et al., 2005; Steven et al., 2014), we note that stronger b-values become increasingly sensitive to shorter molecular distances and the diffusion-weighted signal decay deviates from the monoexponential decay predicted by the Gaussian DTI model after a b-value of (*b* > 1500*s/mm*^2^). That is, the diffusion-weighted signal attenuation *log*(*S*(*b*)/*S*_0_) approximately follows a linear decay up to *b* = 1500*s/mm*^2^. We utilize this observation to adjust for differences in b-values (for 500 < *b* < 1500) between the two sites. Specifically, we estimate the signal for one of the sites at a common harmonized b-value using a linear scaling of the signal in the log-domain. Mathematically, the diffusion signal at a new b-value can be estimated using: *S*=*S*_0_*exp*(–*b_harm_* D̂)where *S* is the diffusion signal, *S*_0_ is the baseline and 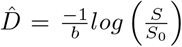 is the diffusion coefficient, and *b* is the original b-value. *b_harm_* is the new b-value of the harmonized data, which is a parameter of choice and we set it to 1000 for all subjects and for both sites in this work (see Table 1, bottom row). For harmonizing b-values greater than 1500 *s/mm*^2^, one could use any of the compressed sensing methods described in (Rathi et al., 2014; Ning et al., 2015b; Fick et al., 2016; Ning et al., 2017, 2015a).

Next, we upsample each diffusion weighted (DW) volume using a 7^*th*^-order B-spline which is shown to perform better than other interpolation schemes (Dyrby et al., 2014). In this study, the harmonized data is resampled to 1.5*mm*^3^ isotropic spatial resolution, which is also a parameter of choice. Next, we use a recently proposed unringing method (Kellner et al., 2015) to remove Gibbs ringing artifacts from each diffusion weighted volume.

#### 2.3.2. Rotation Invariant Spherical Harmonics features

We represent the dMRI signal **S** in a basis of spherical harmonics (SH): **S** *≈* Σ*_l_* Σ*_m_ C_lm_Y_lm_*, where *Y_lm_* are the spherical harmonic basis functions of order *l* and degree *m* with coefficients given by *C_lm_*. From this SH representation, several rotation invariant spherical harmonic (RISH) features at each voxel can be computed as follows (Mirzaalian et al., 2015):

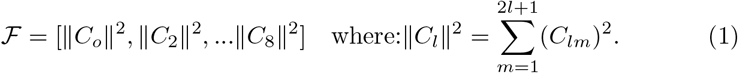

These RISH features can be appropriately scaled to modify the dMRI signal without changing the principal diffusion directions of the fibers (Mirzaalian et al., 2016). Thus, our goal is to estimate a voxel-wise linear mapping of the RISH features between the reference and target sites using matched healthy controls, which can then be used to harmonize the rest of subjects in the target site. We note that this mapping is linear in the SH domain, but non-linear in the dMRI signal domain.

Five RISH feature maps 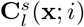 for SH orders of *l* = {0, 2, 4, 6, 8} are computed at each voxel location **x** = (*x, y, z*) ∈ ℝ^3^ for each scanner *s* as follows:

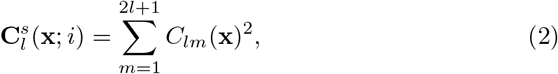

where *i* is the subject number.

#### 2.3.3. Multi-variate template construction using training subjects

Using target scanner RISH features as input, our goal is to learn a voxel-wise linear mapping between the target scanner and the reference scanner. To achieve this, first, the RISH features in the training set are used to create a multi-modal RISH feature template (antsMultiVariateTemplateConstruction (Avants et al., 2010)). Once the template space is constructed separately for each shell (in case of multiple b-value data), we define the expected value of the voxel-wise RISH features as the sample mean 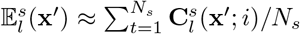 over the number of training subjects *N_s_*, where *s* is the target site or reference site and **x**′ is the voxel location in the template space. Next, we compute voxel-wise linear (scaling only) maps between RISH features of target site (*tar*) and reference site (*ref*) data in the template space using:

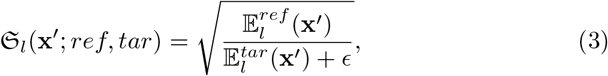

where *l* is the order of the RISH feature and ϵ is a small non-zero constant. Figure 3 shows five mean templates of RISH features for *l* = {0, 2, 4, 6, 8} from left to right for PNC (top row) and BWH (middle row) sites. Note that, each RISH feature captures different frequencies of the dMRI signal. For instance, RISH feature for *l* = 0 captures isotropic components of the diffusion signal, while *l* = 2 is similar to FA and *l* ≥ 4 captures higher order frequencies. Consequently, RISH feature for each *l* represents different microstructural tissue properties of the dMRI signal, which can be modified to harmonize the dMRI data from different sites without changing the underlying fiber orientations and hence the fiber connectivity of the subjects. Note the sharp differences in the RISH feature maps between the two sites, indicating regional and tissue specific non-linear differences between the sites. Figure 3 also shows the scale maps learned for each RISH feature from the training subjects at both sites. As expected, the difference between sites is region and tissue specific.

#### 2.3.4. Harmonization

We apply the linear map (for each RISH feature separately) learned from the training data set to all new subjects in the target site by non-rigid spatial transformation of the linear maps to the native subject space. The non-rigid transformation is obtained by registering the RISH features of each subject to the template space. The inverse of this transformation is applied to the estimated inter-site linear map. The harmonized dMRI signal is then calculated by scaling the SH coefficients of the signal at each voxel in the subject space as follows:

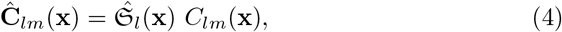

where 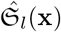 is the scale map in the subject space and 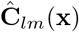 is the scaled SH coefficients. The final diffusion signal is then computed using:

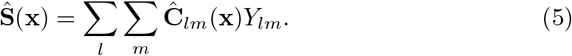

### 2.4. Training set size

In this section we investigate the effect of the size of training subjects on the estimated RISH feature map between sites. We begin by selecting a matched set of subjects at both sites varying in size from 2 to 20 (consecutive even numbers). For each training set size, we generated multiple bootstrap samples of size 100 to estimate the distribution of the scanner differences. The subjects were matched across sites for age, gender and IQ to the best possible extent across sites for each bootstrap sample.

To demonstrate the effect of training data size, in Figure 4, we plot the number of training subjects versus the estimated whole brain mean and standard deviation (std) of the scale map. Our goal is to determine the minimum training set size after which the mean and standard deviation of the scale maps changes minimally, i.e. adding more subjects to the training data does not affect the scale maps. In Figure 4 we show the mean and standard deviation curves for RISH features for order 0, 2 and 4 (order 6 and 8 behave very similar to order 4) separately. For each training size, the mean and std of scale maps is computed in whole brain in 100 bootstrap samples. We observe that the curves become almost stable after a training size of 16, which implies that at-least 16 well matched subjects at each site are needed to learn a robust mapping between sites for dMRI data harmonization. Further, we also observe that the average difference of the mean and std between training size of 18 and 20 is ≤ 0.01. In the rest of this work, we set our training data set size to 20 which can provide robust learning of scanner differences between sites.

**Figure 4:**
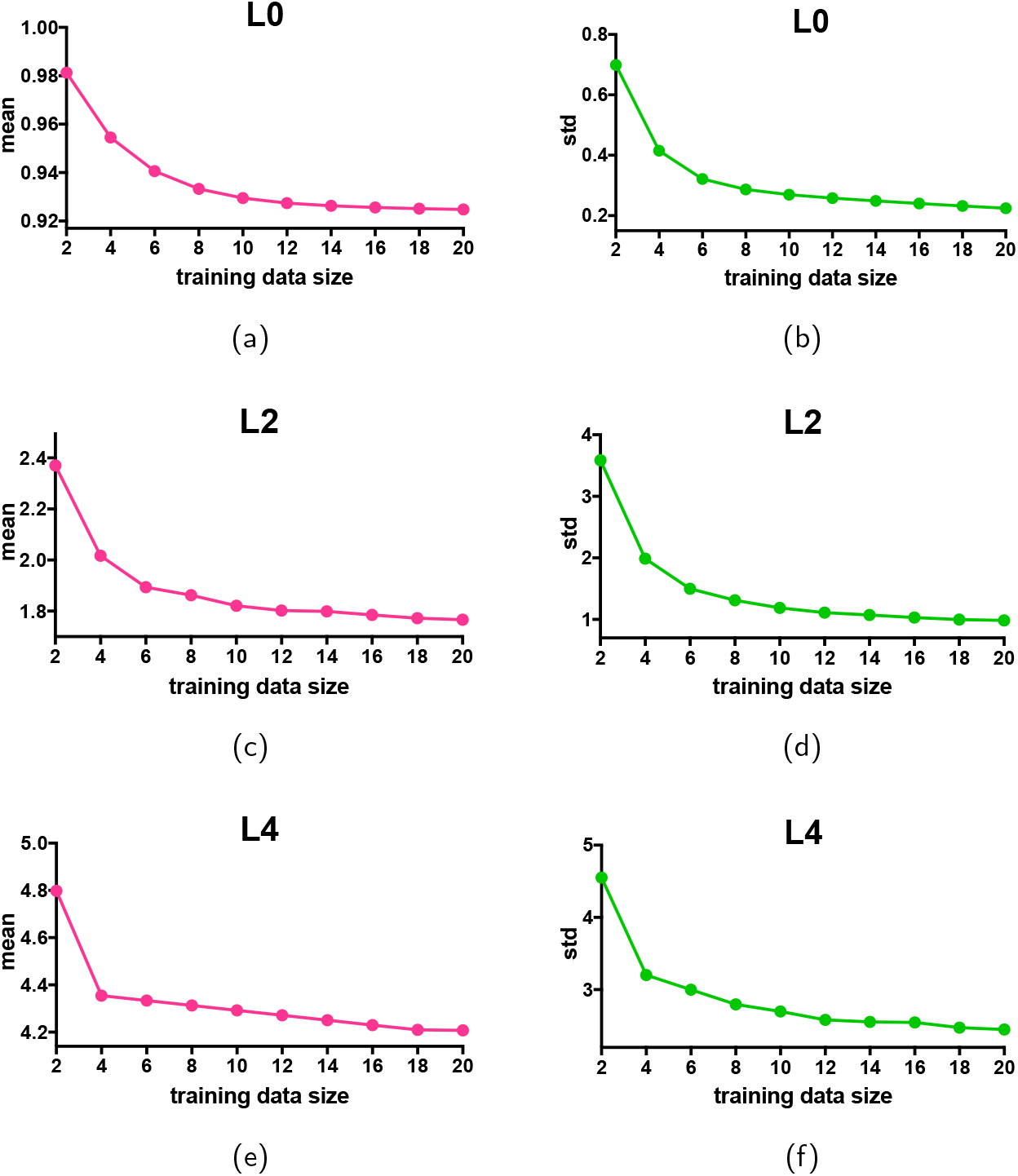
Training size of subjects from 2 to 20 (consecutive even numbers) vs mean (left column-pink) and std (right column-green) of scale maps for RISH features of L0 (top-row), L2 (middle-row) and L4 (bottom-row) to decide how many training subjects are needed to learn the scanner differences across sites. For each training set size, we generate multiple bootstrap samples of size 100. The subjects were matched across sites for age and IQ to the best possible extent across sites for each bootstrap sample.

To provide a more region-specific view, in Figure 5, we depict the differences between the scale maps with a training size of 20 (as “gold standard”) and some representative training data sets of size 2, 12, 16 and 18 for each RISH feature (L0, L2 and L4). Even though we observe large differences between the data sets with 20 subjects and 2 subjects, we see that the voxel-wise differences significantly decrease and the difference maps become more similar after a training size of 16.

**Figure 5:**
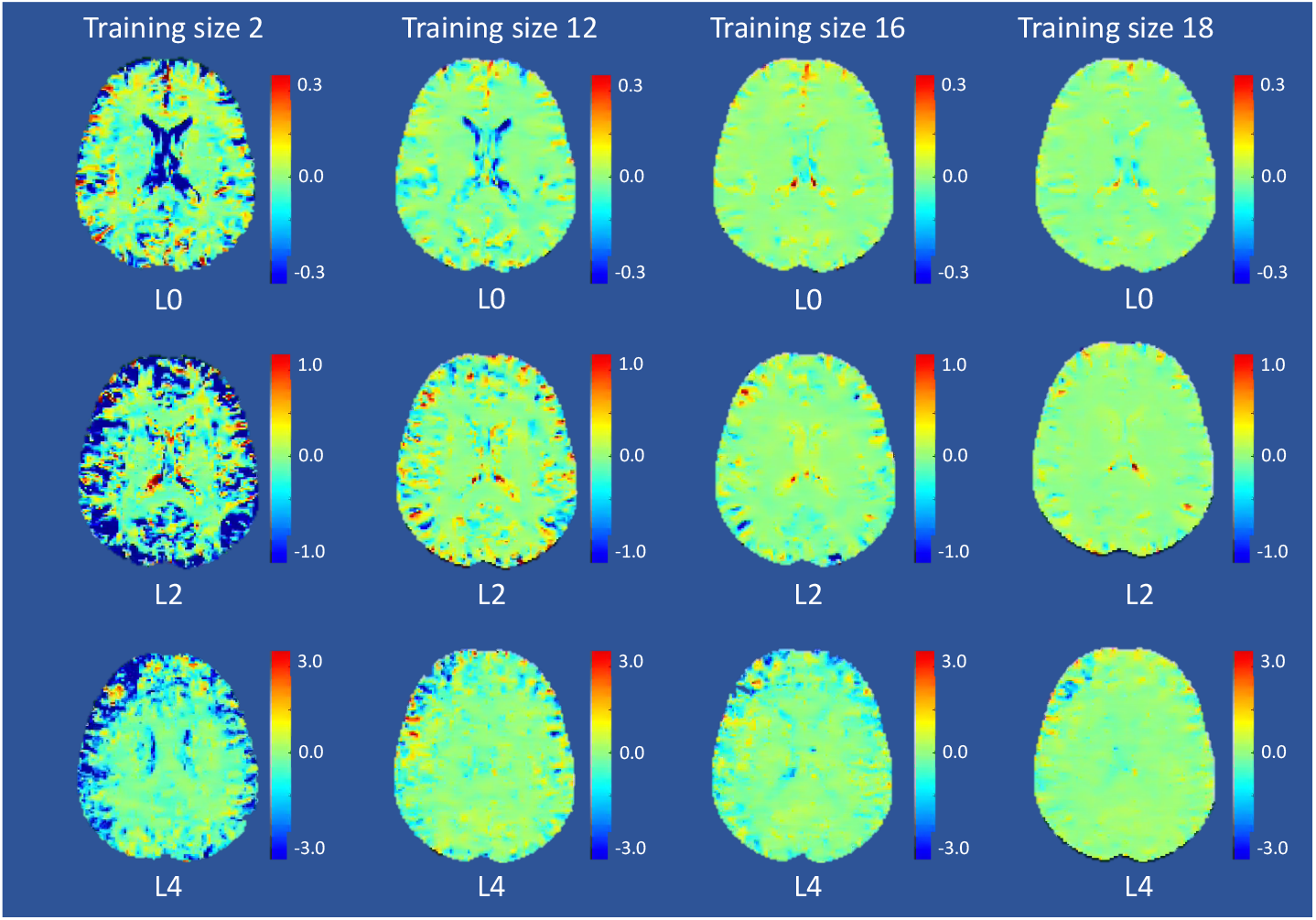
Voxel-wise scale map differences between RISH feature scale map (L0, L2 and L4) estimated with a training size of 20 (as “gold standard”) and some representative training data size of 2, 12, 16 and 18 shown in each column respectively.

## 3. Experiments and Results

### 3.1. Experimental setup

In this section, we describe experiments to evaluate the performance of the proposed algorithm. First, to show that the harmonization works equally well irrespective of the choice of the reference site, we will evaluate the performance of our method using two experiments. In the first experiment, we choose BWH as the reference site and PNC as the target site, whereas in the second experiment we use PNC as the reference site and BWH as the target site. Another aim of these experiments is also to demonstrate the robustness of the proposed technique to preserve group differences despite the size of the test data sets. For example, we evaluate age-related group differences in the PNC data which has a large number of subjects, as well as the BWH site which has a small sample size.

We use dMRI-derived measures of FA, MD and generalized-FA (GFA), which are typically used in neuroimaging studies to understand the effect of harmonization. These measures were also chosen as they are known to change with age in a nonlinear fashion (Yeatman et al., 2014; Lebel et al., 2008) and show different maturational pattern between males and females (Gur et al., 1999; Asato et al., 2010). Hence, our experiments consisted of evaluating the effect sizes between three/two groups (separated by age or sex) before and after harmonization. To investigate performance of the harmonization in different regions of the brain that are known to mature at different speed and be sex dependent, we used the Illinois Institute of Technology (IIT) Human Brain Atlas (Varentsova et al., 2014; Zhang and Arfanakis, 2018). We used 16 different white matter bundles^1^ from this atlas to evaluate the performance of our method. We set a threshold to 0.2 for all subjects to clearly define the regions-of-interest (ROIs) in the IIT probabilistic atlas. Mean FA, MD and GFA were computed in each region for all subjects before and after harmonization to use in the upcoming experiments.

For each of these experiments, we selected 20 right-handed subjects (10 males, 10 females) from each site, matched on age and IQ as described in Section 2.4. The demographic information details about these training subjects was given in Table 2.

In Section 3.2.1, we test the learning (mapping) capabilities and performance of our algorithm on 20 training subjects selected from each site. In Section 3.2.2, we show how the aging and gender effects are preserved after harmonization in a large number of test subjects. In Section 3.2.3, we demonstrate that the proposed harmonization procedure preserves fiber orientation by comparing fiber bundle tracing results before and after harmonization.

### 3.2. Results

#### 3.2.1. Evaluation on training subjects

In Figure 6, we show the mean FA (a, b), mean MD (c, d) and mean GFA (e, f) values in the reference site (red), the target site (green) and the harmonized results (blue) for each of the major white matter bundles (from the IIT atlas) on the training data. In (a, c, e), respectively, we depict the results for FA, MD and GFA with PNC as the reference site and BWH as the target site. In (b, d, f), the experiment is repeated with BWH as the reference site and PNC as the target site. We also observe that the site differences are not uniform but vary in a highly nonlinear fashion across the brain and for all measures. We note that the site differences appear to be more for MD as compared to FA and GFA, which was also reported in (Vollmar et al., 2010).

**Figure 6:**
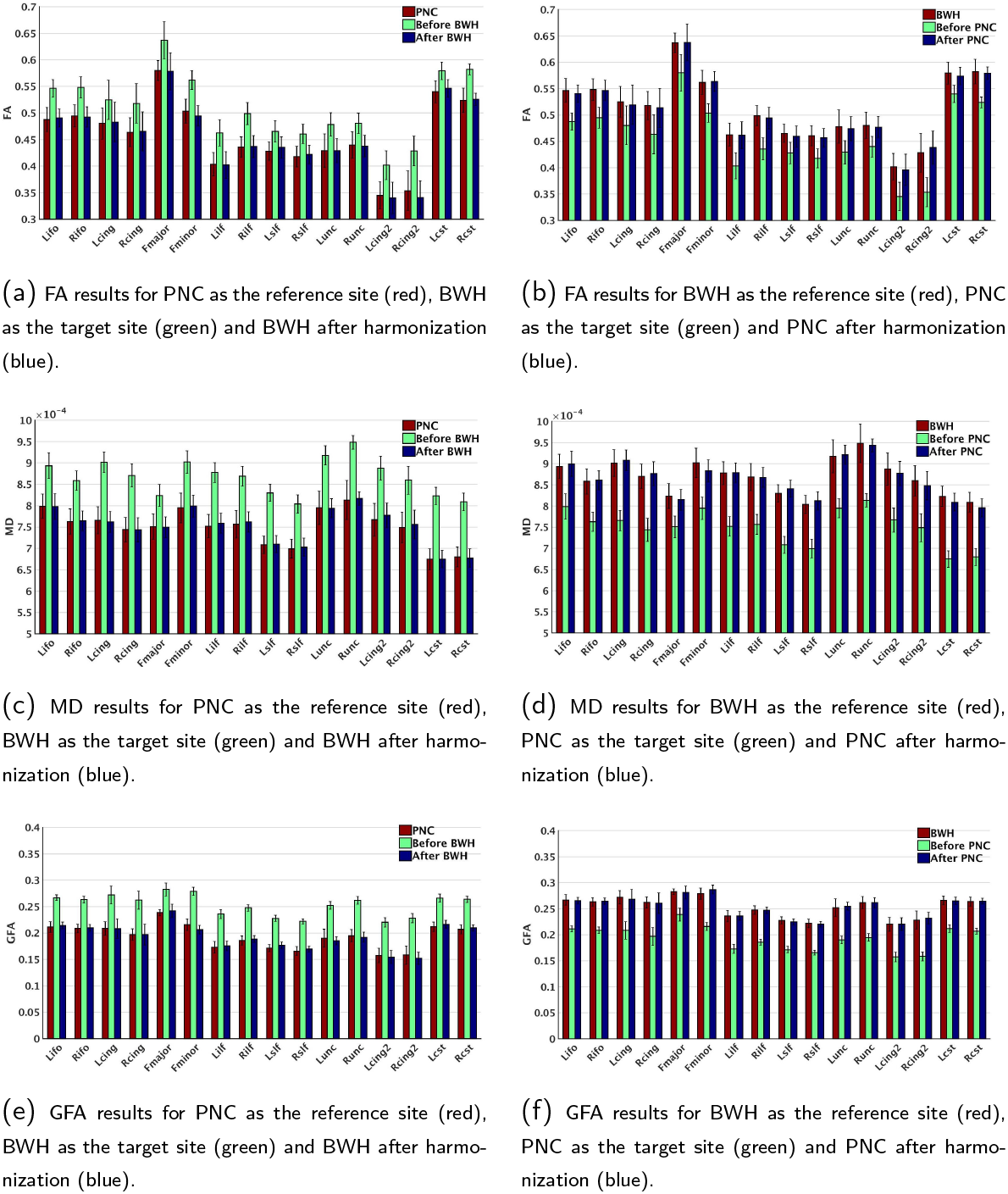
Comparison of diffusion measures (FA, MD and GFA) across sites: red: reference site, green: target site, and blue: after harmonization of target to reference. Left column: PNC is selected as reference site and BWH is selected as target site. Right column: BWH is selected as reference site and PNC is selected as target site. In both scenarios, large statistical differences are observed prior to harmonization. By harmonization, the scanner effects are removed.

To statistically analyze each diffusion measure before and after harmonization, the parametric paired t-test was applied to all major bundles between two sites: (i) reference site and target site (before harmonization); (ii) reference site and harmonized site (after harmonization). See Table 4 for the statistics of PNC as the target site and see Table 5 for the statistics of BWH as the target site. We observe a significant difference between the two sites for all measures (*p* < 1*e* − 4 for all bundles and measures) before harmonization. After harmonization, the statistical difference between controls from both sites is removed for all bundles and measures.

**Table 3:**
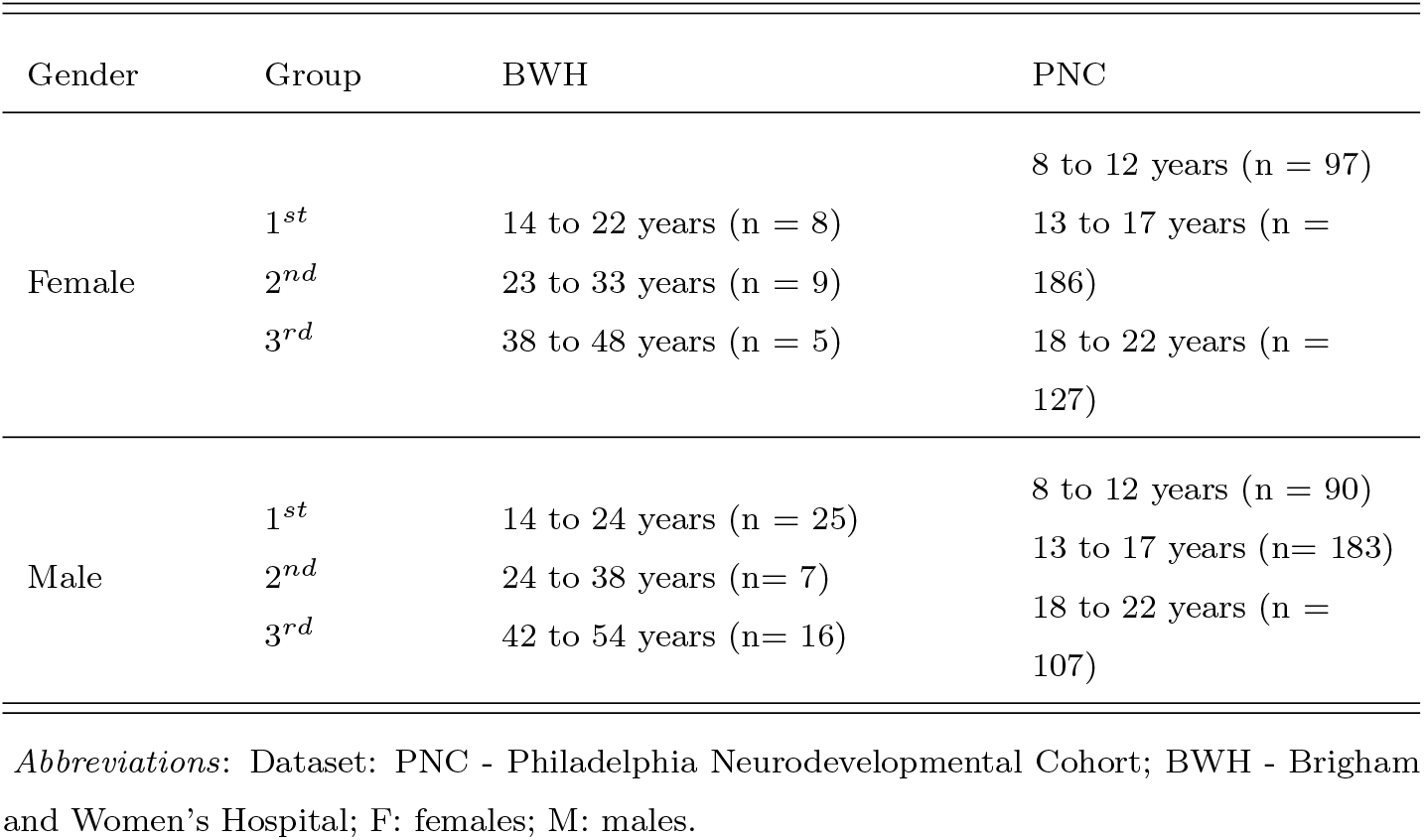
Three age groups in BWH and PNC data for females and males separately.

**Table 4:** The statistics for PNC data: paired t-test was applied to each diffusion measure of all major bundles between two sites: (i) BWH as the reference site and PNC as the target site (the statistics are reported in before PNC rows); (ii) BWH as the reference site and harmonized PNC results (the statistics are reported in after PNC rows).

| Data | P value | t, df | Mean of differences | SD of differences | 95% CI | R<sup>2</sup> | r |
| --- | --- | --- | --- | --- | --- | --- | --- |
| Before PNC:<br>FA | <0.0001 | t=18.14<br>df=15 | -0.04869 | 0.01074 | -0.05442 to<br>-0.04297 | 0.9564 | 0.9861 |
| Before PNC:<br>MD | <0.0001 | t=24.89<br>df=15 | -1.165e-4 | 1.871e-4 | -1.264e-4<br>to 1.065e-4 | 0.9764 | 0.8981 |
| Before PNC:<br>GFA | <0.0001 | t=37.59<br>df=15 | -0.05979 | 0.006362 | -0.06318 to<br>-0.0564 | 0.9895 | 0.9659 |
| After PNC:<br>FA | 0.1825 | t=1.398<br>df=15 | -0.001756 | 0.005025 | -0.004434<br>to<br>0.0009215 | 0.1153 | 0.9968 |
| After PNC:<br>MD | 0.1661 | t=1.456<br>df=15 | -2.708e-6 | 7.442e-6 | -6.674e-6<br>to 1.257e-6 | 0.1238 | 0.9838 |
| After PNC:<br>GFA | 0.7964 | t=0.2626<br>df=15 | 0.0001812 | 0.002761 | -0.00129 to<br>0.001652 | 0.004576 | 0.9913 |

CI: confidence interval; r: correlation coefficient (to observe how effective the pairing is); R<sup>2</sup>: partial eta squared (to observe how big the difference is).

**Table 5:** The statistics for BWH data: paired t-test is applied to each diffusion measure of all major bundles between two sites: (i) PNC as the reference site and BWH as the target site (the statistics are reported in before BWH rows); (ii) PNC as the reference site and harmonized BWH results (the statistics are reported in after BWH rows).

| Data | P value | t, df | Mean of differences | SD of differences | 95% CI | R<sup>2</sup> | r |
| --- | --- | --- | --- | --- | --- | --- | --- |
| Before BWH:<br>FA | <0.0001 | t=18.77<br>df=15 | 0.05251 | 0.01119 | 0.04655 to<br>0.05847 | 0.9591 | 0.9887 |
| Before BWH:<br>MD | <0.0001 | t=24.89<br>df=15 | 1.165e-4 | 1.871e-5 | 1.065e-4 to<br>1.264e-4 | 0.9764 | 0.8981 |
| Before BWH:<br>GFA | <0.0001 | t=37.59<br>df=15 | 0.05982 | 0.006364 | 0.05642 to<br>0.06321 | 0.9895 | 0.9658 |
| After BWH:<br>FA | 0.8119 | t=0.2423<br>df=15 | -0.00075 | 0.01238 | -0.007348<br>to 0.005848 | 0.003898 | 0.9885 |
| After BWH:<br>MD | 0.9235 | t=0.09761<br>df=15 | 2.956e-07 | 1.211e-5 | -6.159e-006<br>to<br>6.751e-006 | 6.348e-4 | 0.9629 |
| After BWH:<br>GFA | 0.7913 | t=0.2694<br>df=15 | 0.0002844 | 0.004222 | -0.001965<br>to 0.002534 | 0.004815 | 0.9847 |

CI: confidence interval; r: correlation coefficient (to observe how effective the pairing is); R<sup>2</sup>: partial eta squared (to observe how big the difference is); ns: not-significant.

#### 3.2.2. Effect size comparison in test subjects

Once a mapping between the sites is estimated from the 20 training subjects (per site), it is applied to the rest of the data set from the target site (i.e., data from all subjects of the target site are updated or harmonized). Any harmonization technique should preserve the inter-subject biological variability and group differences at each site, while only removing scanner related effects. This can be tested by ensuring that the effect sizes between groups is maintained before and after harmonization. White matter maturation (as measured by FA) with age has been well-documented in the literature (Lebel et al., 2008), along with the differential trajectory of this maturation between males and females (Gur et al., 1999). We use this as a test-bed to demonstrate that the effect sizes between groups before and after harmonization is maintained. Specifically, we calculate the effect sizes between groups categorized by age and sex as described in Table 3.

In our first experiment, we calculate the group differences between males and females in FA for each of the three age groups (i.e., matched for age). Our goal is to test if the effect sizes observed in the original test data are preserved after harmonization to a target site. In our second experiment, we calculate the effect sizes due to age before and after harmonization. For both of the experiments, we set: (1) BWH as the reference site and PNC as the target site (see Figure Appendix A.1(a) to see the maturation curves in PNC data); (2) PNC as the reference site and BWH as the target site (see Figure Appendix A.2(a) to see the maturation curves in BWH data).

##### 3.2.2.1. Sex differences (effect sizes) before and after harmonization

We compute the effect sizes using Cohen’s *d* between females and males matched for age for each of the three age groups from Table 3. Mathematically, Cohen’s *d* can be written as: 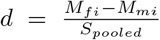, where *M* is the mean FA of the *i^th^* group, *f* represents females, *m* represents males. *S_pooled_* is given by 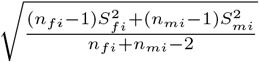 where n is the number of subjects and *S_mi_*; *S_fi_* are the standard deviations for the male and female groups respectively.

###### BWH reference site

In Figure 7(a), we show plots for white matter bundles before and after harmonization. Here BWH is the reference site and PNC is the target site. As can be seen, the effect sizes between the sexes before and after harmonization are almost the same for all age groups, that is, if the effect sizes are small before harmonization, they stay small after harmonization as well. Similar observations can be made for medium and large effect sizes. We however note that, in general, the effect sizes after harmonization are slightly lower than the original, potentially because of some smoothing effects that occur due to interpolation. Nevertheless, these differences are minor and do not change the outcome of statistical analysis.

**Figure 7:**
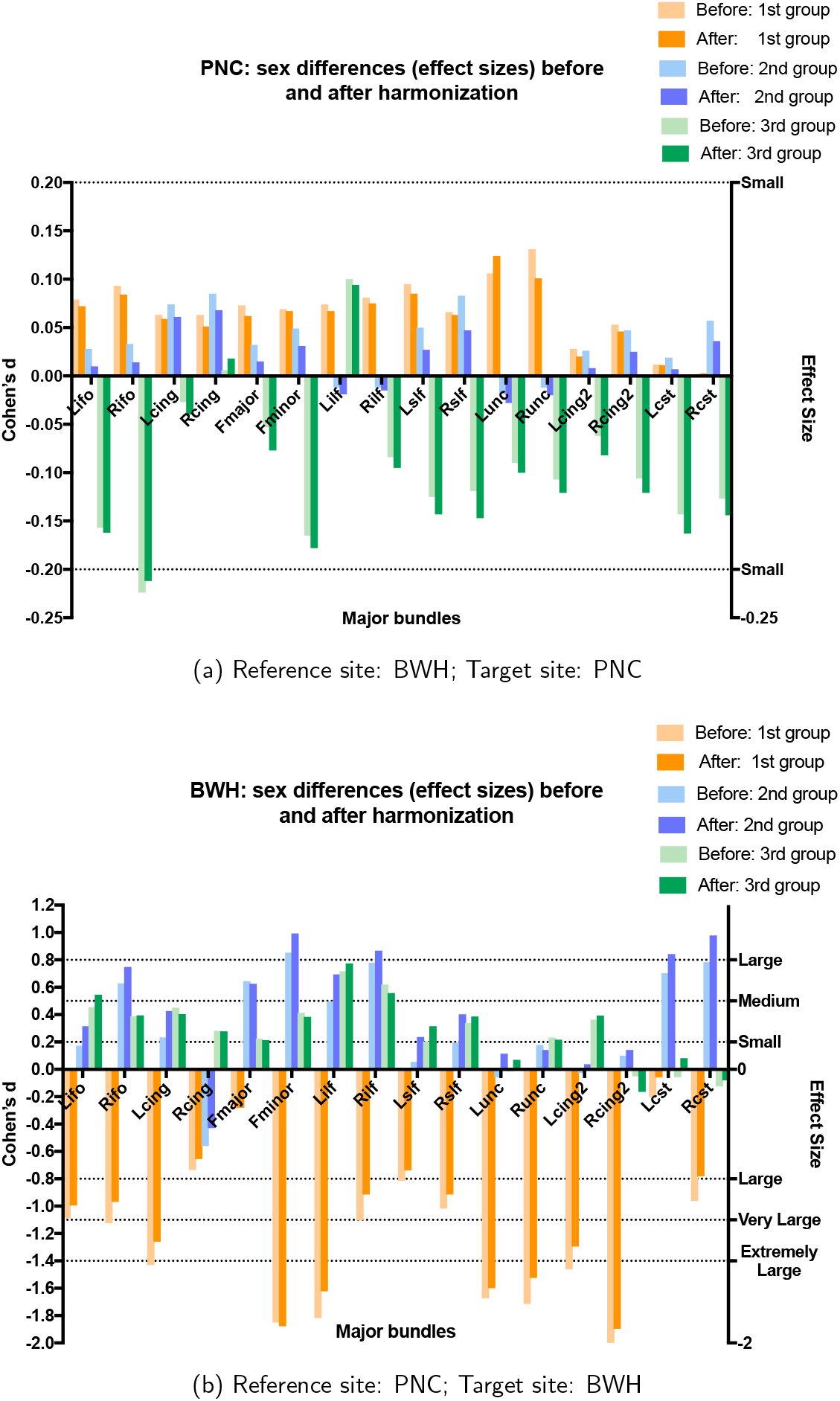
The effect sizes (Cohen’s d) between the sexes for all age groups before and after harmonization. Note that the effect sizes are maintained by the harmonization procedure.

In Table B.1, we provide quantitative values for the effect sizes between groups for BWH as the reference site and PNC as the target site for each major bundle before and after harmonization. We also report the absolute differences (∆) between the effect sizes before and after harmonization. Also reported are results when the effect sizes are grouped into small (d*~*0.2), medium (d*~*0.5), large (d*∼*0.8), very large (d*~*1.1) and extremely large (d*~*1.4) effect sizes. We report the average absolute differences in the effect sizes in each group (Table 6-cyan rows). As can be seen, the effect sizes are preserved after harmonization (i.e., absolute differences in effect sizes before and after harmonization are always close to the original with the average difference being 0.0132).

**Table 6:** Grouped (from small to extremely large) average sex differences in terms of effect sizes before and after harmonization. Cyan rows: the results for BWH as reference and PNC as the target site; Gray rows: the results for PNC as reference and BWH as the target site. ∆ is the absolute difference between effect sizes (before and after harmonization); - implies none of the fiber bundles demonstrated those effect sizes.

|  | *Small* | *Medium* | *Large* | *Very*<br>*Large* | *Extremely*<br>*Large* |
| --- | --- | --- | --- | --- | --- |
| Before PNC | 0.068 | 0.224 | - | - | - |
| After PNC | 0.066 | 0.212 | - | - | - |
| $\Delta$ PNC | 0.002 | 0.012 | - | - | - |
| Before BWH | 0.102 | 0.336 | 0.686 | 0.947 | 1.584 |
| After BWH | 0.154 | 0.379 | 0.720 | 0.884 | 1.440 |
| $\Delta$ BWH | 0.052 | 0.043 | 0.034 | 0.063 | 0.144 |

###### PNC reference site

We also perform a similar analysis for PNC as the reference site and BWH as the target site. At the BWH site, the number of female subjects is very small. Despite this small sample size, the harmonization algorithm preserves the maturation trends very accurately (i.e., trends are very similar to that before harmonization), demonstrating the robustness of the proposed method. However, as seen in Figure 7(b), (and Figure Appendix A.2), small sample sizes can provide misleading (and potentially inaccurate) results as has been shown by several works in the literature. Here, we show these results only to demonstrate that the inter-subject biological variability is preserved by our harmonization algorithm despite the small sample size (test samples) used. We note that no other inferences about sexual dimorphisms can be made from these results from the BWH site.

In Table B.2, we provide quantitative values for the effect sizes for BWH samples before and after harmonization, and their absolute differences for each major bundle. Due to smaller data size of the females and a totally different age range of females and males in each group, unlike the previous experiment, we also observe medium, large, very large and extremely large effect sizes prior to harmonization which are preserved after harmonization (∆ is always < 0.2). Grouping the fiber bundles based on their effect sizes, we once again observe that the effect sizes are preserved after harmonization (Table 6-gray rows), i.e., effect sizes that were small, medium, or large stay small, medium and large respectively after harmonization.

##### 3.2.2.2. Age related effect sizes before and after harmonization

In this experiment, our aim is to show that the effect sizes due to aging are preserved after harmonization. For this purpose, we compute the effect sizes (Cohen’s *d*) between the first and the third age group from Table 3 for both males and females separately. Cohen’s *d* is calculated in a similar fashion as above.

###### BWH reference site

In this case, BWH is the reference site and all data analysis before and after harmonization is done on the PNC site. Since FA increases in young adolescent subjects during maturation (Lebel et al., 2008), it is natural to observe mostly large and positive effect sizes due to aging. Besides, the effect sizes are highly sensitive to gender (see Figure 8). As can be seen, the effect sizes stay almost the same after harmonization in all experiments. In Table B.3, we report the effect sizes of the first and the third age group before and after harmonization and their absolute differences ∆ for males and females separately. Group differences as measured by effect sizes, which are significantly different before harmonization for all bundles, still stay significantly different after harmonization (∆ is always < 0.2). Additionally, the grouped effect size results stay similar after harmonization (Table 7-cyan rows).

**Figure 8:**
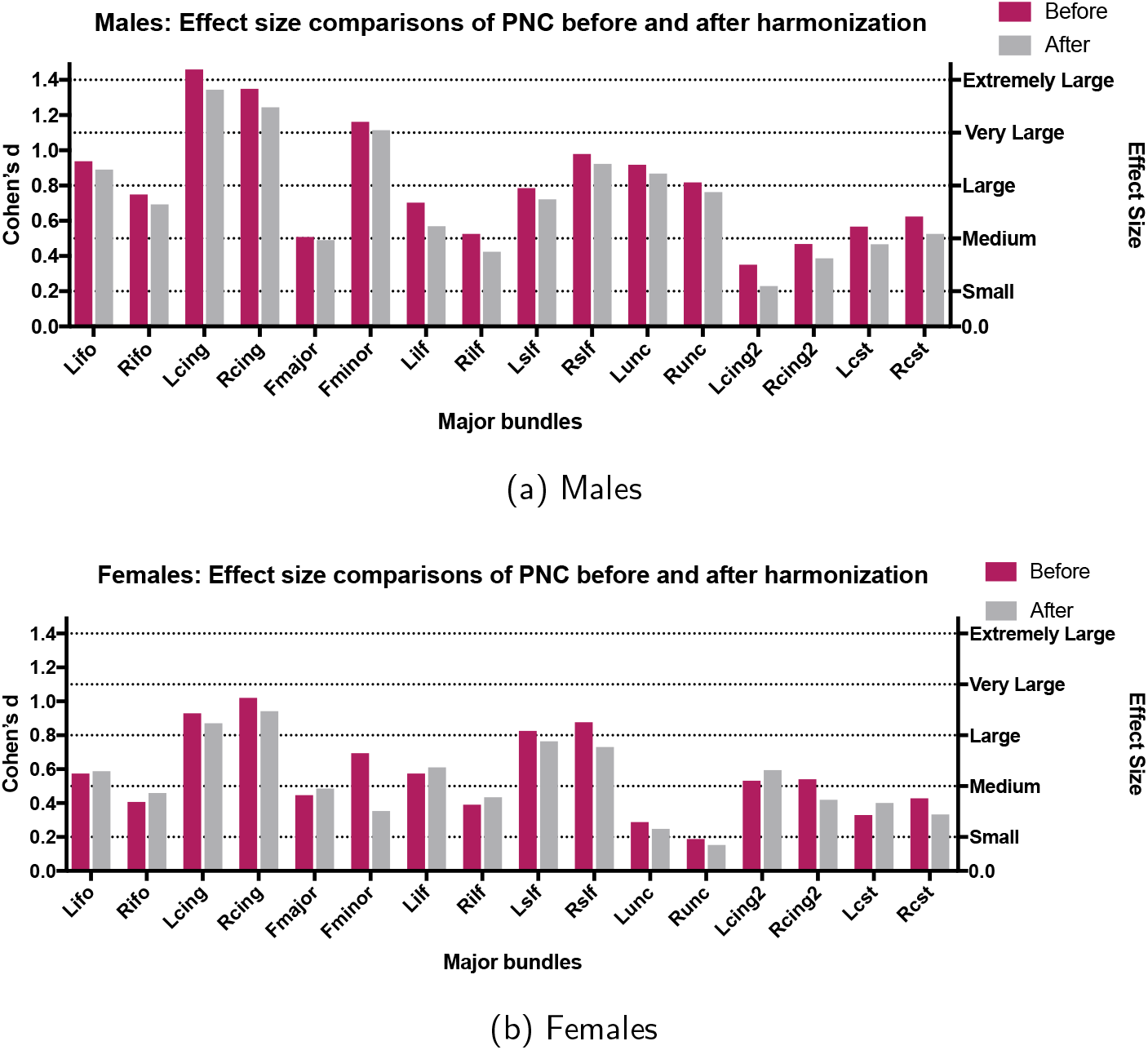
Results for age-related differences between groups with BWH as the reference site and PNC as the target site. The effect sizes (Cohen’s d) between the first and the last group (see Table 3 for the age distribution of the groups) are shown for each gender separately (before harmonization (purple) and after harmonization (gray)).

**Table 7:** Comparison of the average grouped (from small to extremely large) aging effect sizes before and after harmonization. Cyan rows: the results for BWH is reference and PNC is the target site; Gray rows: the results for PNC is reference and BWH is the target site. ∆ is the absolute difference between effect sizes (before and after harmonization); - means their is no value for the corresponding effect size.

|  | *Small* | *Medium* | *Large* | *Very*<br>*Large* | *Extremely*<br>*Large* |
| --- | --- | --- | --- | --- | --- |
| Before PNC | 0.188 | 0.389 | 0.615 | 0.913 | 1.324 |
| After PNC | 0.153 | 0.372 | 0.538 | 0.844 | 1.235 |
| $\Delta$ PNC | 0.035 | 0.017 | 0.077 | 0.069 | 0.089 |
| Before BWH | 0.092 | 0.315 | 0.722 | 0.924 | 1.381 |
| After BWH | 0.090 | 0.283 | 0.660 | 0.822 | 1.338 |
| $\Delta$ BWH | 0.002 | 0.032 | 0.062 | 0.102 | 0.043 |

In this regard, we would also like to point the results of age-dependent maturation curves in the PNC data set. As can be seen in Figure Appendix A.1, the maturation curves are accurately preserved by the harmonization algorithm. When PNC data is the target site (i.e., PNC data is updated for harmonization), we see a robust trend in maturation of different white matter bundles consistent with those reported in the literature (Paus et al., 2001; Paus, 2010).

###### PNC reference site

We also perform a similar analysis for PNC as the reference site and BWH as the target site (i.e., BWH data was harmonized and analyzed before and after harmonization). Due to small sample size and differences in age-ranges, the maturation curves and the effect sizes do not match with those from the much larger PNC data set. However, to clarify once more, our aim is only to validate the harmonization performance regardless of the underlying trends in the data. As can be seen, the harmonization procedure preserves the trends as well as the effect sizes. In Table B.4, we report the effect sizes and ∆ for the BWH site before and after harmonization for males and females respectively. The effect sizes are preserved after harmonization for all white matter bundles (see also Figure 9) and for each group (Table 7-gray rows).

**Figure 9:**
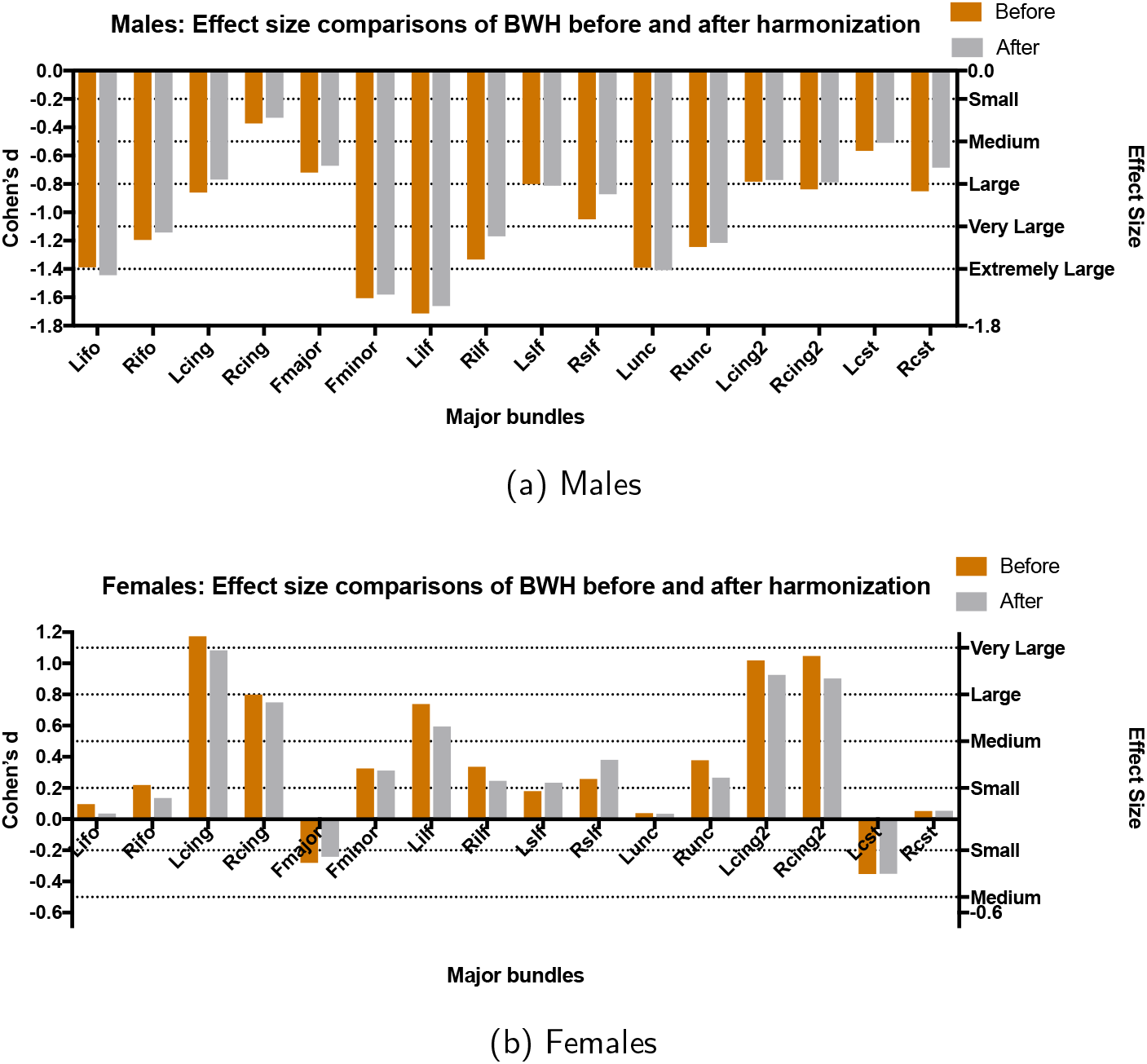
Results for age-related differences between groups with PNC as the reference site and BWH as the target site. The effect sizes (Cohen’s d) between the first and the last group (see Table 3 for the age distribution of the groups) are shown for each gender separately (before harmonization (orange) and after harmonization (gray)).

#### 3.2.3. Tractography analysis

In order to ensure that our harmonization method (which involves modifying the dMRI signal) does not in any way to change the fiber orientations, we performed whole brain tractography using a multi-tensor unscented Kalman filter (UKF) method (Malcolm et al., 2010; Reddy and Rathi, 2016). The same parameters were used to generate whole brain tracts from the original and harmonized dMRI data. Next, the White Matter Query Language was utilized (WMQL) (Wassermann et al., 2016) to extract specific anatomical white matter bundles from the whole brain tracts. Figure 10 depicts WMQL results for corticospinal tract (CST) and the inferior occipital-frontal fibers (IOFF) before (blue) and after (pink) harmonization. After extracting the tracts from a subject before and after harmonization, the Bhattacharyya overlap distance (*B*) was used to quantify the overlap between the tracts (Rathi et al., 2013):

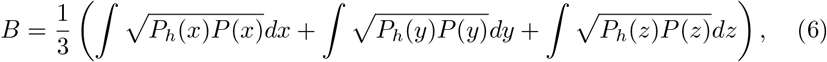

where *P*(.) represents the ground truth spatial probability distribution of the fiber bundle, *P_h_*(.) is the spatial probability distribution of the tracts from the harmonized data and (*x*, *y*, *z*) ∈ ℝ^3^ are the fiber coordinates. *B* is 1 for a perfect match between two fiber bundles and 0 for no overlap at all. We observed very high overlap greater than 0.93 for all fiber bundles indicating that fiber orientation is well preserved by the harmonization algorithm.

**Figure 10:**
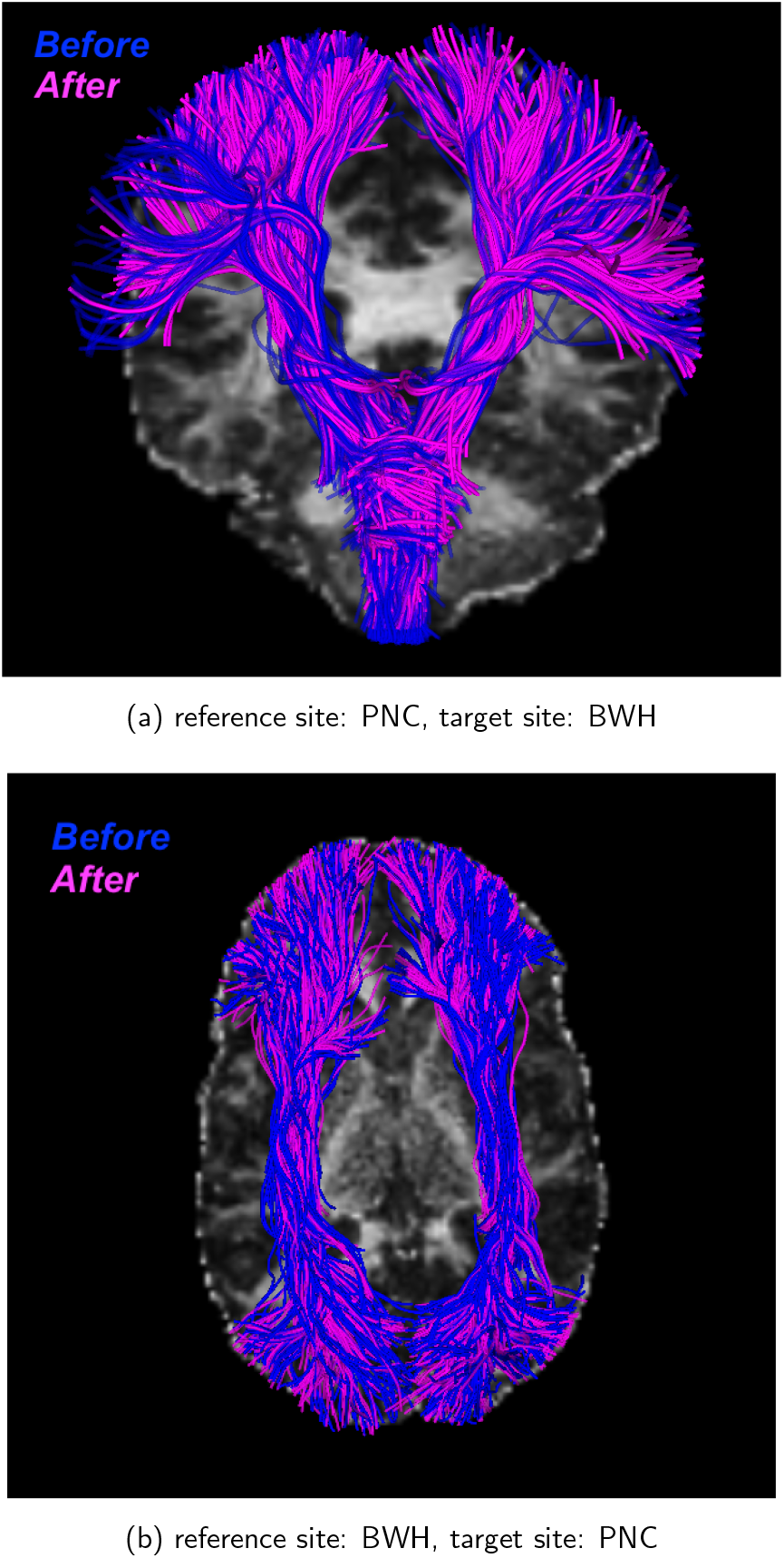
Significant (> 93%) overlap is seen in CST and IOFF fiber bundles before and after harmonization. Blue: before harmonization; Pink: after harmonization.

## 4. Discussion and Conclusion

We believe that accurate harmonization of dMRI data is of utmost importance to allow for a large-scale data-driven way to understand brain disorders. In this paper, we presented a harmonization method to retrospectively remove scanner-specific differences from the raw dMRI signal across various sites, even if acquired with different acquisition parameters. The harmonization procedure requires a well-matched set of controls across sites to learn the mapping between sites.

Acquisition parameters, magnetic field inhomogeneities, coil sensitivity, and other scanner related effects can cause non-linear changes in the signal in different tissue types. To remove these site effects, we first mapped the b-values from each site to a canonical b-value of 1000*s/mm*^2^ and resampled the data to 1.5^3^*mm*^3^ (Section 2.3.1). Later, we utilized RISH features that are able to capture different frequency components of the diffusion signal to learn the inter-site differences (Section 2.3.2). In Figure 3, we showed that the scanner related differences are substantially different for sub-cortical gray, versus the neighboring white matter region or the distant cortical gray matter regions. Further, these differences can be captured selectively by the different frequency bands of the SH basis (i.e., in different RISH features).

We note that, the methodology proposed here harmonizes the raw dMRI signal in a model-independent manner. Further, dMRI data harmonization has to be done only once. Thus, any subsequent analysis will necessarily be consistent, unlike methods that work with model-specific measures such as FA, which are obtained at the last stage of the processing pipeline. Note that, it is not clear how non-linear scanner effects affect the downstream processing and model fitting of dMRI data. Consequently, we recommend that dMRI data be harmonized at the earliest possible processing stage.

Using several experiments, in this paper, we evaluated our method’s performance on two independent sites: PNC with 800 healthy controls and BWH with 70 healthy controls. Our results lead us to conclude the following: (i) At-least 16 to 18 well matched healthy controls from each site were required to learn a robust mapping that can capture only site-related differences. (ii) Irrespective of the effect size (small, medium or large), the proposed harmonization procedure preserved the effect sizes after harmonization. (iii) The harmonization procedure also ensured that the fiber orientation directions were left unchanged.

In this paper, we investigated a method to harmonize dMRI data retrospectively when traveling subjects are not available. Scanner-specific effects from multiple sites can be best captured by acquiring data in quick succession from a set of traveling human subjects. In this case, the scanner specific differences can be obtained from these traveling subjects and subsequently used for data harmonization, and the learned difference mapping could be applied to the unseen subjects in multi-site studies. Evaluating our algorithm on multi-site data from traveling human subjects will form part of our future study.

## 5. Acknowledgements

The authors would like to acknowledge the following grants which supported this work: R01MH102377, (PI: Dr. Marek Kubicki), R01MH097979 (PI: Dr. Yogesh Rathi).

## Appendix A Age-related trends in FA, before and after harmonization

**Figure Appendix A.1.**
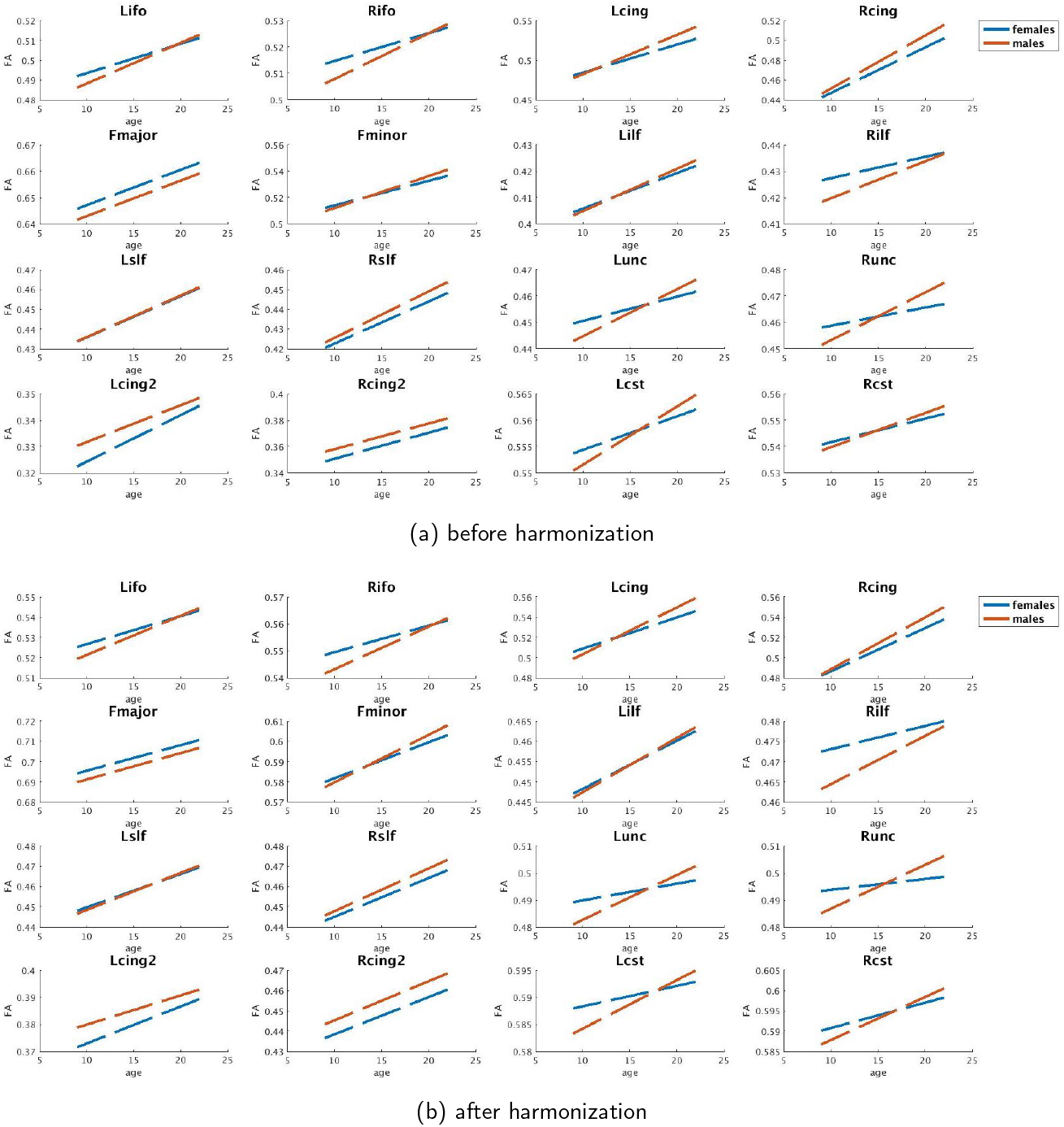
Reference Site: BWH, before and after harmonization female (blue) and male (orange) age vs FA curves of PNC for each major white matter bundle.

**Figure Appendix A.2.**
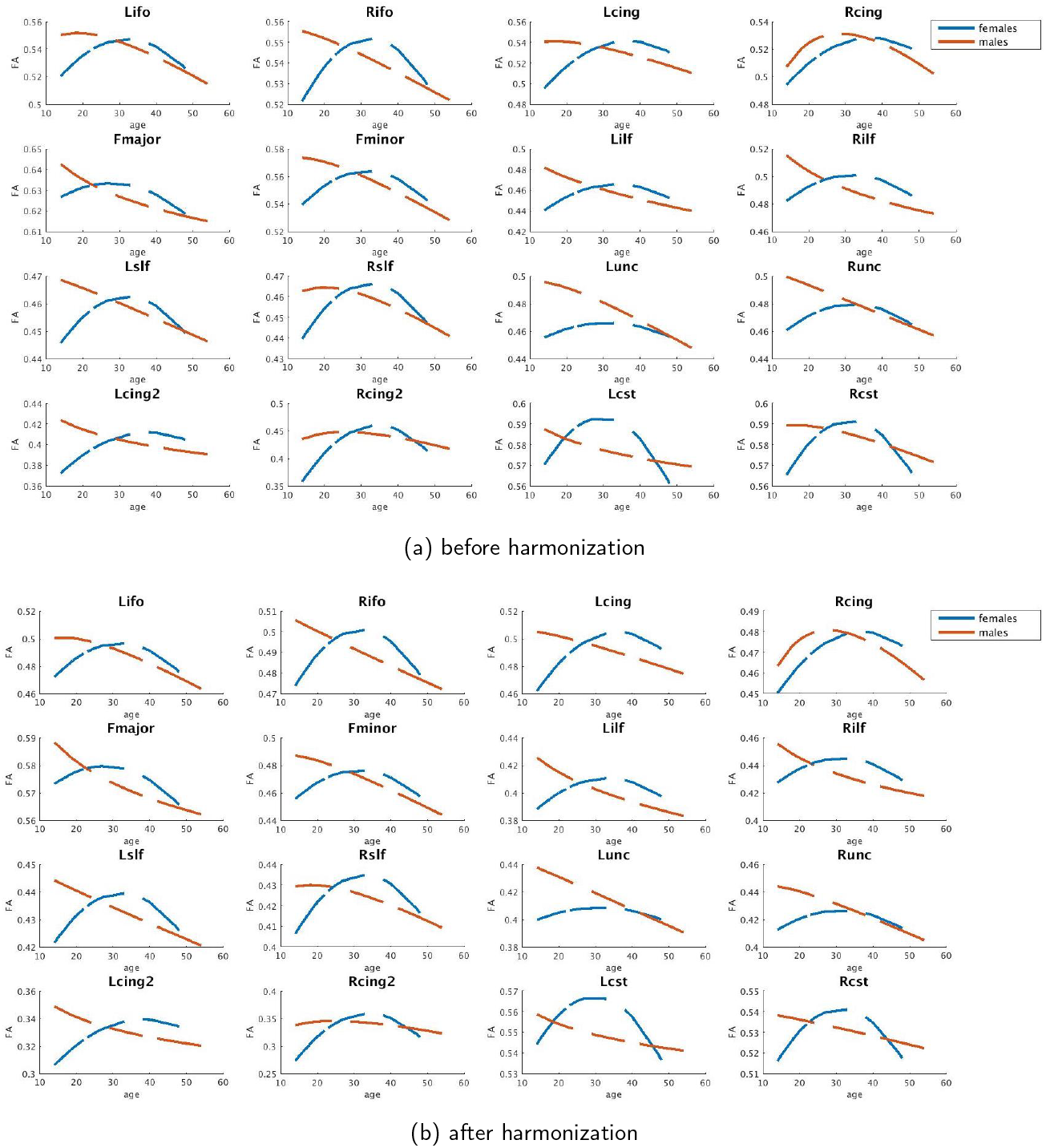
Reference Site: PNC, before and after harmonization female (blue) and male (orange) age vs FA curves of BWH for each major white matter bundle.

## Appendix B Effect sizes before and after harmonization for BWH as the reference site and PNC as the target site

### Appendix B.1. Analysis of sex differences

**Table B.1:**
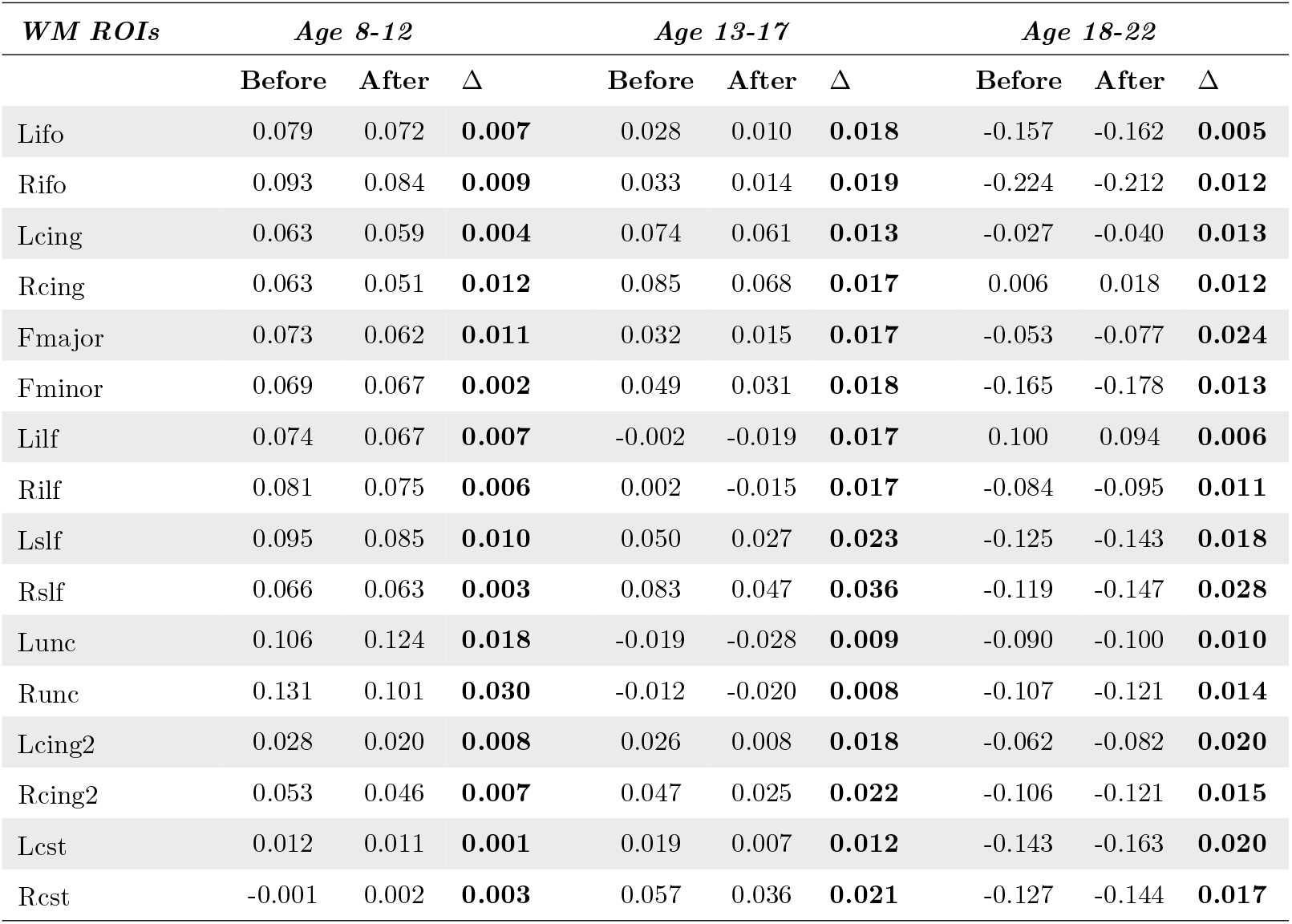
Target site: PNC. Sexual dimorphism effect sizes effect sizes before and after harmonization for each major bundle. Absolute differences (∆) between before and after harmonization effect sizes are observed to be *<* 0.2 in all cases.

**Table B.2:**
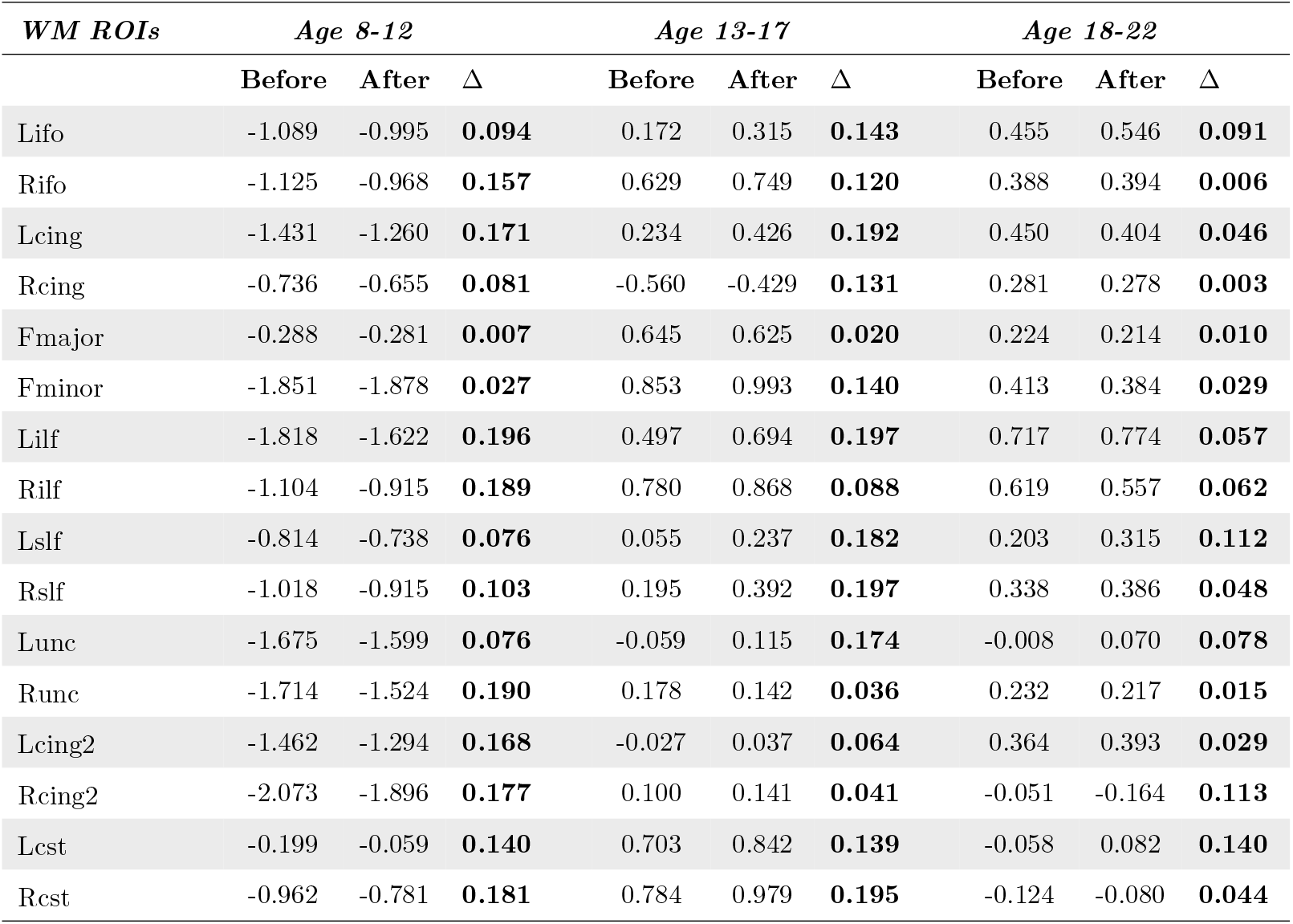
Target site: BWH. Sexual dimorphism effect sizes before and after harmonization for each major bundle. Absolute differences (∆) between before and after harmonization effect sizes are observed to be *<* 0.2 in all cases.

| *WM ROIs* | *Age 8-12* |  |  | *Age 13-17* |  |  | *Age 18-22* |  |  |
| --- | --- | --- | --- | --- | --- | --- | --- | --- | --- |
| | Before | After | $\Delta$ | Before | After | $\Delta$ | Before | After | $\Delta$ |
| Lifo | -1.089 | -0.995 | **0.094** | 0.172 | 0.315 | **0.143** | 0.455 | 0.546 | **0.091** |
| Rifo | -1.125 | -0.968 | **0.157** | 0.629 | 0.749 | **0.120** | 0.388 | 0.394 | **0.006** |
| Lcing | -1.431 | -1.260 | **0.171** | 0.234 | 0.426 | **0.192** | 0.450 | 0.404 | **0.046** |
| Rcing | -0.736 | -0.655 | **0.081** | -0.560 | -0.429 | **0.131** | 0.281 | 0.278 | **0.003** |
| Fmajor | -0.288 | -0.281 | **0.007** | 0.645 | 0.625 | **0.020** | 0.224 | 0.214 | **0.010** |
| Fminor | -1.851 | -1.878 | **0.027** | 0.853 | 0.993 | **0.140** | 0.413 | 0.384 | **0.029** |
| Lilf | -1.818 | -1.622 | **0.196** | 0.497 | 0.694 | **0.197** | 0.717 | 0.774 | **0.057** |
| Rilf | -1.104 | -0.915 | **0.189** | 0.780 | 0.868 | **0.088** | 0.619 | 0.557 | **0.062** |
| Lslf | -0.814 | -0.738 | **0.076** | 0.055 | 0.237 | **0.182** | 0.203 | 0.315 | **0.112** |
| Rslf | -1.018 | -0.915 | **0.103** | 0.195 | 0.392 | **0.197** | 0.338 | 0.386 | **0.048** |
| Lunc | -1.675 | -1.599 | **0.076** | -0.059 | 0.115 | **0.174** | -0.008 | 0.070 | **0.078** |
| Runc | -1.714 | -1.524 | **0.190** | 0.178 | 0.142 | **0.036** | 0.232 | 0.217 | **0.015** |
| Lcing2 | -1.462 | -1.294 | **0.168** | -0.027 | 0.037 | **0.064** | 0.364 | 0.393 | **0.029** |
| Rcing2 | -2.073 | -1.896 | **0.177** | 0.100 | 0.141 | **0.041** | -0.051 | -0.164 | **0.113** |
| Lcst | -0.199 | -0.059 | **0.140** | 0.703 | 0.842 | **0.139** | -0.058 | 0.082 | **0.140** |
| Rcst | -0.962 | -0.781 | **0.181** | 0.784 | 0.979 | **0.195** | -0.124 | -0.080 | **0.044** |

### Appendix B.2. Analysis of aging

**Table B.3:** Target site: PNC. Age related effect sizes before and after harmonization for each major bundle. Absolute differences (∆) between before and after harmonization effect sizes are observed to be *<* 0.2 in all cases.

| *WM ROIs* | *Males* |  |  | *Females* |  |  |
| --- | --- | --- | --- | --- | --- | --- |
| | Before | After | $\Delta$ | Before | After | $\Delta$ |
| Lifo | 0.938 | 0.891 | **0.046** | 0.574 | 0.588 | **0.014** |
| Rifo | 0.749 | 0.693 | **0.056** | 0.407 | 0.459 | **0.052** |
| Lcing | 1.459 | 1.345 | **0.114** | 0.929 | 0.870 | **0.059** |
| Rcing | 1.349 | 1.245 | **0.104** | 1.021 | 0.942 | **0.079** |
| Fmajor | 0.508 | 0.491 | **0.017** | 0.447 | 0.486 | **0.039** |
| Fminor | 1.162 | 1.114 | **0.048** | 0.694 | 0.353 | **0.341** |
| Lilf | 0.703 | 0.570 | **0.133** | 0.575 | 0.610 | **0.035** |
| Rilf | 0.526 | 0.425 | **0.101** | 0.391 | 0.434 | **0.043** |
| Lslf | 0.785 | 0.722 | **0.064** | 0.825 | 0.764 | **0.061** |
| Rslf | 0.979 | 0.923 | **0.056** | 0.877 | 0.731 | **0.146** |
| Lunc | 0.918 | 0.868 | **0.050** | 0.288 | 0.248 | **0.040** |
| Runc | 0.818 | 0.763 | **0.054** | 0.188 | 0.153 | **0.035** |
| Lcing2 | 0.351 | 0.229 | **0.122** | 0.532 | 0.595 | **0.063** |
| Rcing2 | 0.468 | 0.387 | **0.080** | 0.541 | 0.419 | **0.122** |
| Lcst | 0.566 | 0.467 | **0.099** | 0.329 | 0.400 | **0.071** |
| Rcst | 0.625 | 0.526 | **0.099** | 0.428 | 0.333 | **0.095** |

**Table B.4:**
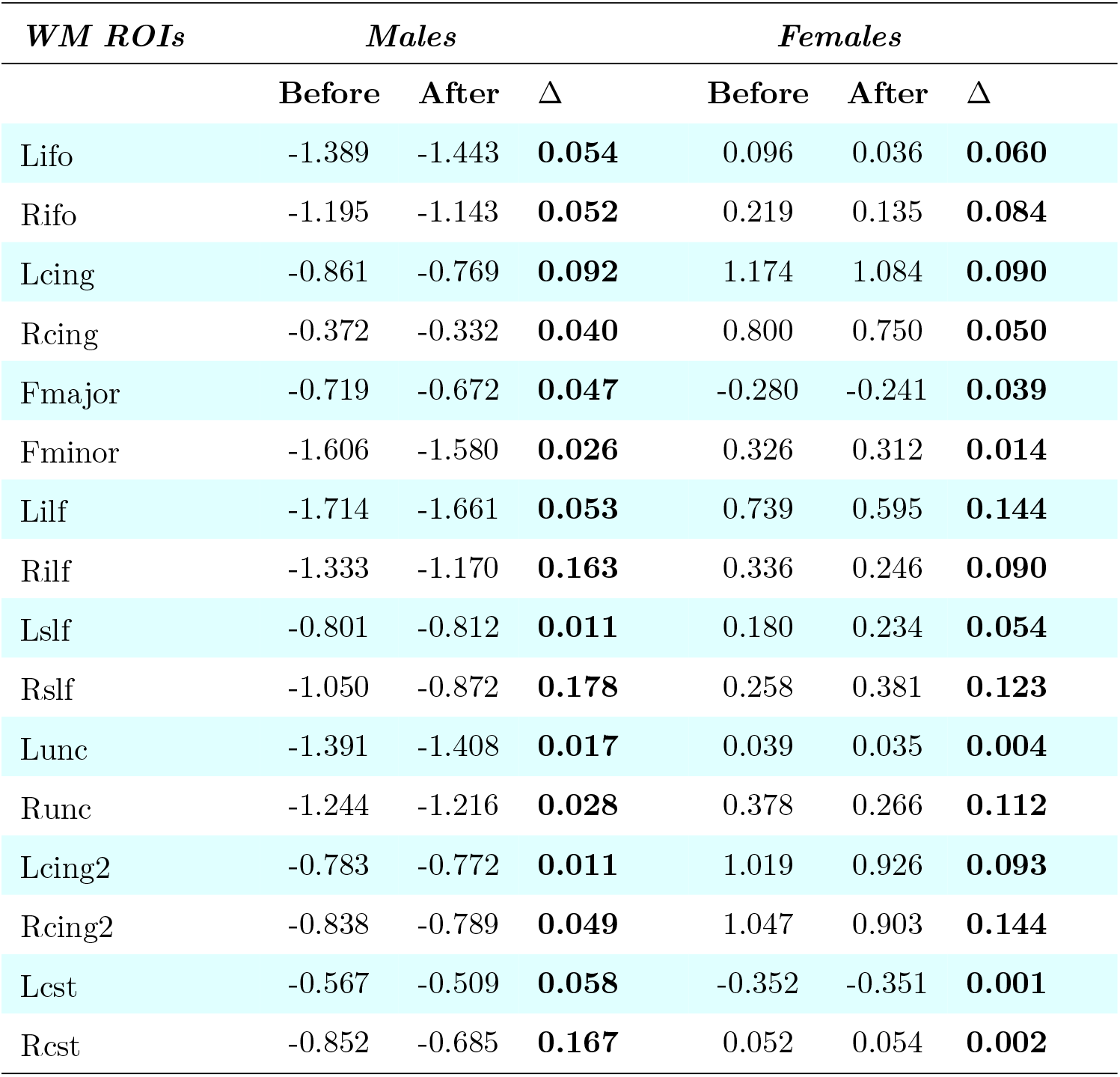
Target site: BWH. Age related effect sizes before and after harmonization for each major bundle. Absolute differences (∆) between before and after harmonization effect sizes are observed to be *<* 0.2 in all cases.

| *WM ROIs* | *Males* |  |  | *Females* |  |  |
| --- | --- | --- | --- | --- | --- | --- |
| | Before | After | $\Delta$ | Before | After | $\Delta$ |
| Lifo | -1.389 | -1.443 | **0.054** | 0.096 | 0.036 | **0.060** |
| Rifo | -1.195 | -1.143 | **0.052** | 0.219 | 0.135 | **0.084** |
| Lcing | -0.861 | -0.769 | **0.092** | 1.174 | 1.084 | **0.090** |
| Rcing | -0.372 | -0.332 | **0.040** | 0.800 | 0.750 | **0.050** |
| Fmajor | -0.719 | -0.672 | **0.047** | -0.280 | -0.241 | **0.039** |
| Fminor | -1.606 | -1.580 | **0.026** | 0.326 | 0.312 | **0.014** |
| Lilf | -1.714 | -1.661 | **0.053** | 0.739 | 0.595 | **0.144** |
| Rilf | -1.333 | -1.170 | **0.163** | 0.336 | 0.246 | **0.090** |
| Lslf | -0.801 | -0.812 | **0.011** | 0.180 | 0.234 | **0.054** |
| Rslf | -1.050 | -0.872 | **0.178** | 0.258 | 0.381 | **0.123** |
| Lunc | -1.391 | -1.408 | **0.017** | 0.039 | 0.035 | **0.004** |
| Runc | -1.244 | -1.216 | **0.028** | 0.378 | 0.266 | **0.112** |
| Lcing2 | -0.783 | -0.772 | **0.011** | 1.019 | 0.926 | **0.093** |
| Rcing2 | -0.838 | -0.789 | **0.049** | 1.047 | 0.903 | **0.144** |
| Lcst | -0.567 | -0.509 | **0.058** | -0.352 | -0.351 | **0.001** |
| Rcst | -0.852 | -0.685 | **0.167** | 0.052 | 0.054 | **0.002** |

## Footnotes

1 Abbreviations: forceps major (Fmajor), forceps minor (Fminor), fornix (Fornix), cingulum (cingulate gyrus portion) (Lcing and Rcing for left and right hemispheres respectively), cingulum (hippocampal portion) (Lcing2 and Rcing2), corticospinal tract (Lcst and Rcst), inferior fronto-occipital fasciculus (Lifo and Rifo), inferior longitudinal fasciculus (Lilf and Rilf), superior longitudinal fasciculus (Lslf and Rslf), uncinate fasciculus (Lunc and Runc)

